# Cancer persister cells activate NMDARs to survive ferroptosis

**DOI:** 10.64898/2026.07.13.738168

**Authors:** Simona Punzi, Ilaria Villanti, Guido Gatti, Davide Cittaro, Gemma Crupi, Michela Prunella, Nicola Altini, Elisa Guerrera, Giulia Casaroli, Giovanni Filippo Marco Gallo, Claudia Felici, Oronza A. Botrugno, Emanuele Tanzi, Vitoantonio Bevilacqua, Antonella Nai, Laura Silvestri, Giovanni Tonon

**Author notes:** These Authors contributed equally to the work (Simona Punzi, Ilaria Villanti).

## Abstract

Upon treatment, cancer cells engage non-genetic adaptations, including tolerance and subsequent persistence, to survive therapy. Eliciting programmed cancer cell death in these persister cells (PCs) remains a primary goal in oncology. We found that ferroptosis is the programmed cell death mechanism most deregulated in persisters by some of the most widely used therapeutic regimens, including platinum-based therapies, which combined with ferroptosis inducers ablate persister colorectal cancer cells. Conversely, persisters emerging from topoisomerase inhibitor regimens withstand ferroptosis and ferroptotic inducers, increasing instead intracellular iron concentration. We found that topoisomerase inhibitors trigger the Xc^-^ antiporter axis (*via* SLC7A11 and CD44) increasing both intracellular cystine, to activate GPX4, and extracellular glutamate. Glutamate then engages the NMDA receptors (NMDARs), which are essential in neurotransmission but recently reported to be deregulated also in cancer cells. In PCs, NMDARs stimulate intracellular Ca^2+^ uptake and trigger the AKT/NFE2L2 axis, thereby engaging a cytoprotective program to cope with oxidative stress. Furthermore, we found that NFE2L2 increases the distance between the endoplasmic reticulum and mitochondria while reducing mitochondrial ROS in PCs. The synergistic inhibition of both the standard (Xc^-^ antiporter) and this novel NMDAR/NFE2L2 axis resensitizes PCs to ferroptosis. These data provide new opportunities to improve the efficacy of widely used therapeutic regimens.

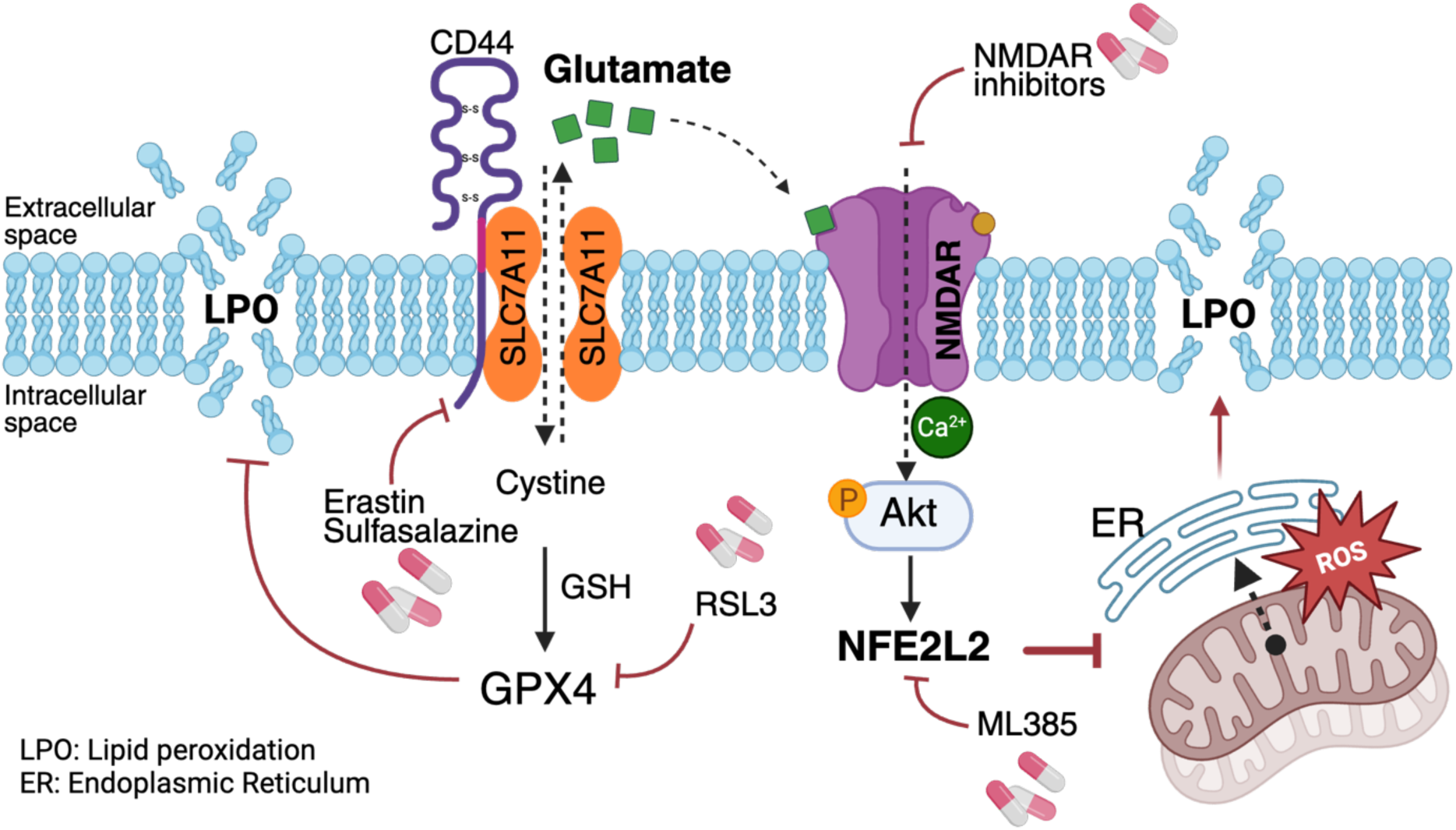

- Persister cells induced by FOLFOX are sensitive to ferroptosis while resistant upon FOLFIRI treatment
- Increased extracellular glutamate activates a NMDAR/NFE2L2 axis
- NFE2L2 copes with oxidative stress by inhibiting juxtaposition between ER and mitochondria
- Inhibition of Xc^-^ antiporter alongside NMDAR/NFE2L2 is required to trigger ferroptosis in FOLFIRI persister cells

## Introduction

Cancer cells exposed to chemotherapy enter in a state of tolerance, a response driven by autophagy that allows them to withstand even exceedingly high drug concentrations^1^. Upon prolonged drug exposure, tolerance morphs into a persistence state, an alternative form of non-genetic adaptation contributing to the emergence of resistance^1^. Persister cells (PCs) are slow-cycling and undergo specific metabolic and epigenetic reprogramming to survive therapy, which includes chromatin remodelling^2^, aldehyde dehydrogenase (ALDH) activity or fatty acid oxidation enhancement^3–4^ and maintenance of mitochondrial fitness through PINK1-dependent mitophagy^1^.

The eradication of PCs remains one of the main aims in cancer therapy^5^. Cells may undergo regulated cell death (RCD) through several pathways, including apoptosis, necroptosis, autophagy, pyroptosis, ferroptosis, and cuproptosis^6^. However, PCs manage to evade these regulated mechanisms. For example, following targeted and conventional cancer therapies, PCs elude both intrinsic and extrinsic pathways of apoptosis^7–8^. As a result, there is growing interest in triggering and exploiting alternative RCD mechanisms to eliminate PCs in cancer^9^.

Ferroptosis is a form of RCD elicited by the accumulation of intracellular free iron, which activates pro-oxidative enzymes, alters mitochondrial activity and tilt the balance from monounsaturated (MUFA) towards polyunsaturated fatty acid (PUFA). The net result is the peroxidation of lipids on the plasma membrane followed by cellular demise^10^. Compounds eliciting ferroptosis have been devised that overcome resistance to traditional therapies and enhance the impact of radiotherapy and immunotherapy^11^. However, cancer cells mitigate the threat of overwhelming membrane lipid peroxidation leveraging the concerted activity of several enzymes and metabolites. These strategies involve various antioxidant defence mechanisms, including cystine-GPX4 (Xc^-^antiporter system), FSP1-CoQ_10_ and GTP-BH2/4 detoxification reactions^12^. Specifically, the Xc^-^antiporter system combines the entrance into the cell of cystine with the ejection of glutamate across the plasma membrane. Cystine then increases the synthesis of glutathione (GSH), a reducing agent used by glutathione peroxidase 4 (GPX4) to convert toxic phospholipid hydroperoxides into non- toxic lipid alcohols, thus protecting the cell from oxidative stress^13^. While high extracellular glutamate inhibits Xc^−^ antiporter and induces ferroptosis^14^, it remains unclear whether it exerts alternative role in controlling ferroptosis, beyond the Xc^-^ system.

Here, we explored which RCD path is triggered in PCs following chemotherapy, which remains the most widely used cancer therapy, with the goal to unravel potential resistance mechanisms implemented by PCs to withstand RCD.

## Results

### FOLFOX- but not FOLFIRI-PCs are susceptible to ferroptosis induction

We focused on colorectal cancer (CRC), a major cancer type with limited therapeutic strategies, where cells enter a persistence state following chemotherapy^1,15–16^. To determine whether CRC PCs are susceptible to any form of RCD after treatment with standard of care, we probed the pathways responsible for the various forms of cell death in PCs derived from the CRC cell line HCT116 treated with the two standard-of care treatments for CRC^17^, FOLFOX (5-fluorouracil + oxaliplatin) or FOLFIRI (5-fluorouracil + SN38). We assessed the variation in gene expression on pathways associated with the different forms of RCD, leveraging our recently published single cell RNA-seq (scRNA-seq) data^1^. With the prominent exception of ferroptosis, we did not detect any deregulation in gene expression of various RCD programs in PCs, including apoptosis, which was also assessed *via* Annexin V labelling (Fig.1 a, Suppl. Fig. 1a-b)^1^. Instead, we found a strong increase in the expression of ferroptosis-related genes (Fig. 1a). Prompted by these findings, we anticipated a strong synergistic response between standard chemotherapy treatments and ferroptosis inducers such as Erastin and RSL3, which respectively inhibit the cystine-glutamate antiporter system Xc^−^ and glutathione peroxidase 4 (GPX4)^18–19^. Both Erastin and RSL3 almost eliminated PCs after FOLFOX treatment (Fig. 1b), suggesting that indeed this treatment is associated with an enhanced, potentially exploitable vulnerability to ferroptosis.

**Figure 1.**
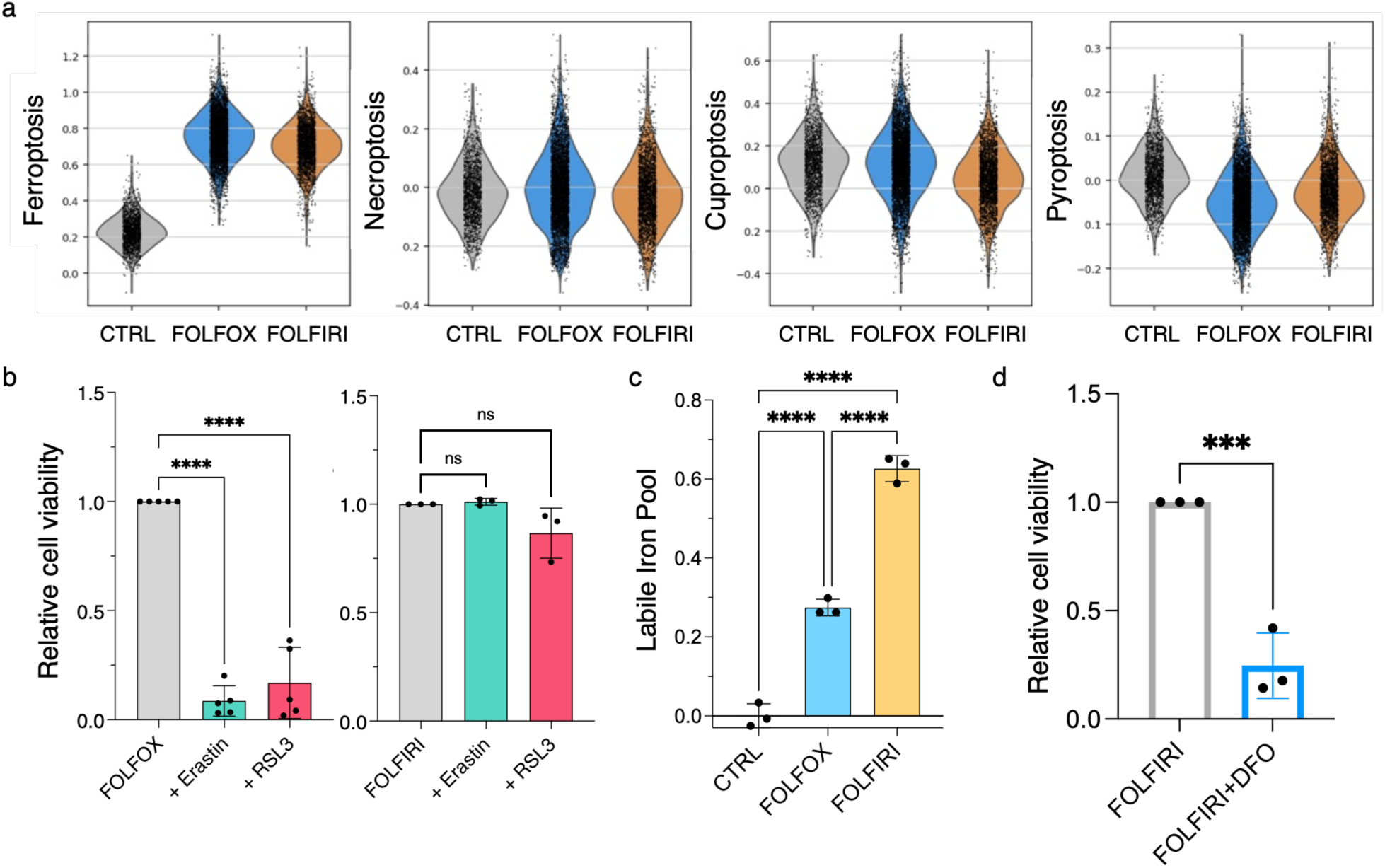
FOLFOX- but not FOLFIRI-PCs are susceptible to ferroptosis induction. **a.** Single cell RNA sequencing analysis of main programmed cell death mechanisms in HCT116 PCs and parental CTRL. **b.** Relative cell viability of HCT116 PCs obtained upon FOLFOX (n=5 biological replicates; mean±SD) or FOLFIRI (n=3 biological replicates; mean±SD) administration and then treated with ferroptosis inducers Erastin and RSL3. Significant differences among groups calculated by one way ANOVA followed by Dunnett’s post-test (****: P<0.0001; ns: not significant). **c.** Labile iron content quantification by flow cytometry analysis upon calcein administration in parental CTRL and FOLFOX- and FOLFIRI-PCs. Significant differences among groups calculated by ANOVA followed by Tukey post-test. Significant differences among groups calculated by one way ANOVA followed by Dunnett’s post-test (n=3 biological replicates; mean±SD; ****: P<0.0001). **d.** Relative cell viability of HCT116 PCs obtained upon FOLFIRI (n=3 biological replicates; mean±SD) and then treated with deferoxamine (DFO). Significant differences among groups calculated by Student’s t-test (***: P<0.001).

Despite a similar increase in ferroptosis-related genes (Fig. 1a), FOLFIRI treatment associated with ferroptosis inducers failed to show any effect (Fig. 1b). Since iron (Fe^2+^) levels are directly linked to the induction of ferroptosis, we anticipated that FOLFIRI treatment would be associated with reduced intracellular Fe^2+^ levels. Surprisingly, however, labile Fe^2+^ content was significantly higher in PCs treated with FOLFIRI than with FOLFOX (Fig. 1c and Suppl. Fig. 1c). Additionally, in FOLFIRI-treated cells, this accumulation of Fe^2+^ was associated with a significantly more pronounced increase in ferritin (FTH1), and curiously in transferrin receptor (TFRC), markers of Fe^2+^ storage and uptake, respectively (Suppl. Fig. 1d), when compared with FOLFOX-treated cells. Taken together, these data demonstrate that FOLFIRI treatment confers resistance to ferroptosis even in the presence of a robust increase in cellular Fe^2+^. This accumulation, likely mediated by deregulated TFRC-driven extracellular transport, highlights a clear uncoupling of Fe^2+^, FTH1, and TFRC within the homeostatic regulatory network^20^.

Previous evidence has suggested that Fe^2+^ may be required for cancer stem-like cells and tumour-initiating cells survival ^21^. Nevertheless, how Fe^2+^ functionally contributes to the maintenance of a treatment-resistant phenotype is still not fully elucidated. We then hypothesized that to withstand FOLFIRI, PCs rely on Fe^2+^ accumulation. FOLFIRI-pretreated cells were challenged with the potent Fe^2+^ chelator deferoxamine (DFO), which acts by sequestering intracellular iron^22^. Treatment with DFO markedly impaired the viability of PCs while promoting an increase in Propidium Iodide (PI)-positive cells (Fig. 1d and Suppl. Fig. 1e), thus validating the hypothesis that Fe^2+^, despite its role in triggering ferroptosis, in FOLFIRI-PCs exerts nonetheless a pro-survival function.

### FOLFIRI hyper-activates the Xc^-^ antiporter system through CD44

During cell death triggered by ferroptosis, Fe^2+^ overload stimulates oxidation of membrane phospholipids, thereby initiating peroxidation^10^. We then determined the levels of lipid peroxidation after both treatments. FOLFIRI-PCs exhibited lipid peroxidation levels comparable to those observed in control cells, in contrast to the substantial increase seen in FOLFOX- PCs (Fig. 2a). This reinforces the notion that FOLFIRI-PCs activate protective mechanisms to prevent the onset of ferroptosis, despite elevated Fe^2+^ levels. The Xc^-^ cystine/glutamate antiporter system plays a central role in inhibiting ferroptosis through the increase of intracellular cystine followed by the synthesis of GSH^13^. We hence explored whether the resistance towards ferroptosis in FOLFIRI-PCs might be related to an increased activity of this system. Indeed, FOLFIRI-PCs showed a robust increase of both NADPH, which provides reducing equivalents to regenerate GSH^23^, and GSH *per se*, which were instead not enhanced in FOLFOX-PCs (Fig. 2b-c). Also GPX4, a powerful antioxidant enzyme belonging to the glutathione peroxidase family^24^, was strongly upregulated in FOLFIRI- but not in FOLFOX-PCs (Fig. 2d), in line with previous observation that Xc^-^-mediated cystine uptake promotes not only GSH, but also GPX4 protein synthesis^25^.

**Figure 2.**
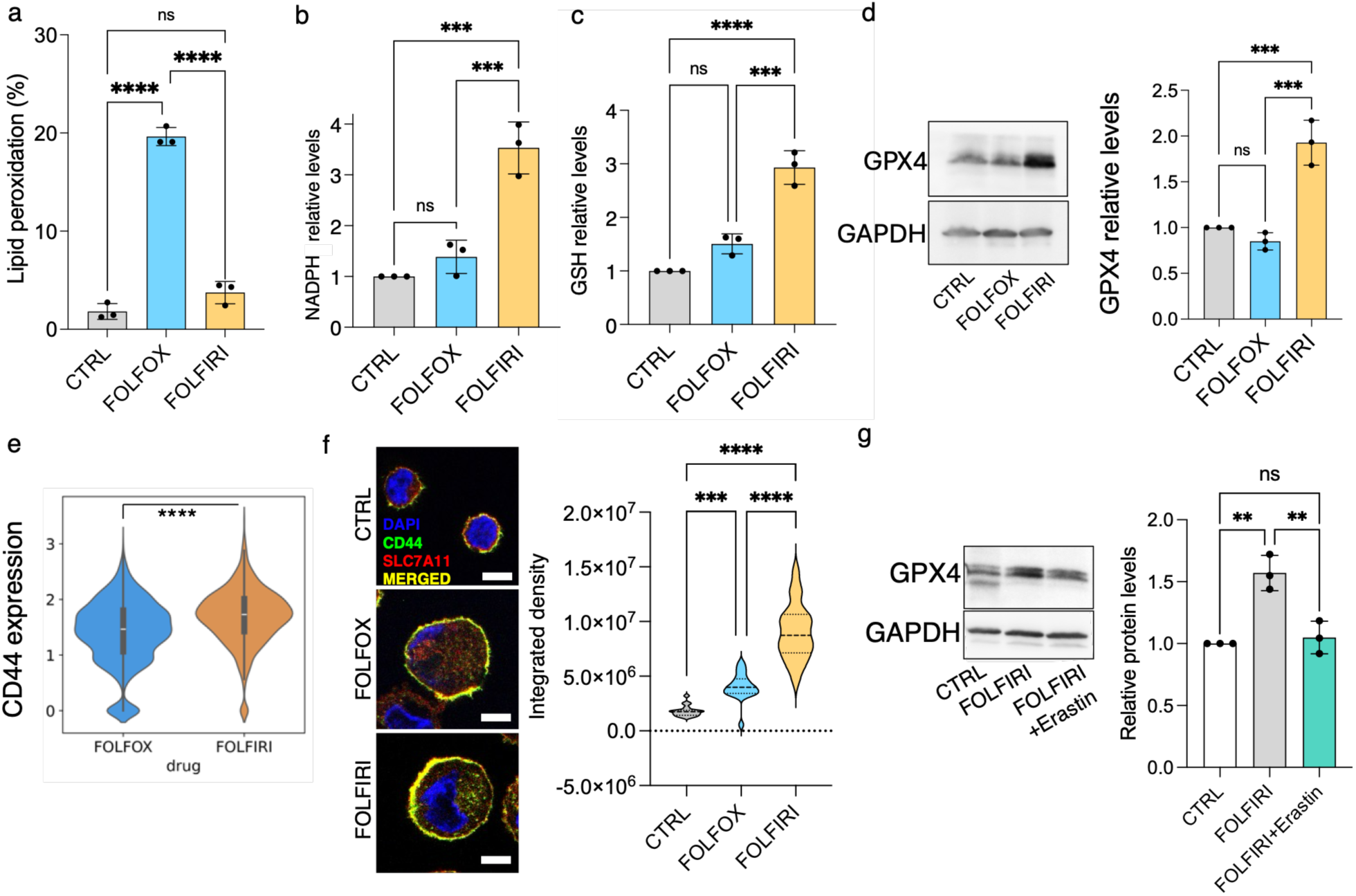
FOLFIRI hyper-activates SCL7A11 axis through CD44 regulation. **a.** Lipid peroxidation levels quantified as percentage positivity (%) in PCs and parental CTRL by flow cytometry analysis. Significant differences among groups calculated by ANOVA followed by Tukey post-test (n=3 biological replicates; mean±SD; ****: P<0.0001; ns: not significant). **b-c.** NADPH (**b**) and Glutathione (GSH) (**c**) cellular content in PCs compared to the parental CTRL. Significant differences among groups calculated by ANOVA followed by Tukey post-test (n=3 biological replicates; mean±SD; ***: P<0.001; ****: P<0.0001). **d.** Western blot analysis of GPX4 levels and its quantification in PCs compared to the parental CTRL. GAPDH was used for normalization. Significant differences among groups calculated by ANOVA followed by Tukey post-test (n=3 biological replicates; mean±SD; ***: P<0.001). **e.** Transcription levels of CD44 in FOLFOX- and FOLFIRI-PCs at single cell level. Significant differences among groups calculated by a Student t-test (****: P<0.0001). **f.** Immunofluorescence analysis probing co-localization of CD44 and SLC7A11 in CTRL, FOLFOX- or FOLFIRI-PCs. Significant differences among groups calculated by ANOVA followed by Tukey post-test (n=3 biological replicates biological replicas and 20 cells analysed; mean±SD; ***: P<0.001; ****: P<0.0001). **g.** Western blot analysis of GPX4 levels and its quantification in FOLFIRI-PCs compared to the parental CTRL and in presence of Erastin. GAPDH was used for normalization. Significant differences among groups calculated by ANOVA followed by Tukey post-test (n=3 biological replicates; mean±SD; **: P<0.01; ns: not significant).

In all, these results suggest that, despite a strong increase in cellular Fe^2+^, FOLFIRI-PCs manage to bypass ferroptosis through the activation of the Xc^-^ antiporter system, leading to the increase of GSH biosynthesis and of GPX4^26^.

Unlike FOLFOX, the FOLFIRI regimen includes SN38, the active metabolite of the topoisomerase inhibitor irinotecan. Treating cancer cells with SN38 alone had similar effects than FOLFIRI (Suppl. Fig. 2a). Furthermore, the co-treatment of SN38 and ferroptosis inducers failed to alter the survival of PCs, consistent with the results obtained using FOLFIRI (Suppl. Fig 2b). SN38 interferes with DNA unwinding and induces DNA double strand breaks. Accordingly, cancer cells resistant to topoisomerase inhibition often exhibit alterations in DNA remodelling and chromatin accessibility^27^. We wanted to determine first whether PCs treated with FOLFIRI demonstrated a different level of chromatin compaction, and next if the expression of specific genes was associated with the treatment. To this end, we exploited single cell GET-Seq (scGET-Seq), a technology that probes both open and closed chromatin, at the single cell level^28^. We have previously shown that upon chemotherapy, chromatin becomes more compact in PCs^1^. However, comparing chromatin status between the two treatments, revealed that chromatin remains more accessible in FOLFIRI-PCs than in FOLFOX-PCs (Suppl. Fig. 2c). We next sought to identify the subset of expressed genes located within these regions specifically open in FOLFIRI- PCs. Integration of scGET-seq with scRNA-seq data showed that CD44, a gene residing within one of these selectively open loci, is also overexpressed (Suppl. Fig. 2d). CD44 is a multifunctional cell surface protein that stabilizes the SLC7A11 subunit of the Xc^-^ antiporter system *via* its chaperone CD98hc at the plasma membrane, leading to increased resistance to ferroptosis^29^. We found that both the transcription and the surface protein levels of CD44 were significantly higher in FOLFIRI- than in FOLFOX-PCs (Fig. 2e and Suppl. Fig. 2e). In addition, confocal microscopy and a proximity ligation assay (PLA) confirmed the colocalization of CD44 and SLC7A11 (Fig. 2f and Suppl. Fig 2f-g). To verify that CD44 was functionally involved in SLC7A11 activity, we measured the change in intracellular cystine uptake in response to FOLFIRI and CD44 knockdown (Suppl. Fig. 2h). FOLFIRI-PCs robustly increased the intake of a cystine analogue, which was abrogated by CD44 silencing (Suppl. Fig. 2i), suggesting that FOLFIRI increases the expression of CD44, which in turn cooperates with SLC7A11 to increase the influx of cystine and activate anti-ferroptosis responses.

As detailed above, FOLFIRI treatment increases both GSH and GPX4 in PCs cells. Curiously, the inhibition of the Xc^-^ antiporter obtained by Erastin, CD44 or SLC7A11 knockdown (Suppl. Fig. 2j-k), as well as the inhibition of GPX4 by RSL3, were all unable to elicit ferroptosis in these cells, unlike in other persistence models^30^. As a matter of fact, the combination of Erastin with FOLFIRI brought back GPX4 to the levels seen in untreated cells (Fig. 2g). This suggests that, in conjunction with GSH and GPX4 increase, additional mechanisms elicit ferroptosis resistance in FOLFIRI-PCs.

### Glutamate activates NMDAR and induces cytoprotection

Since the Xc^-^ antiporter couples cystine uptake with glutamate efflux, we investigated whether glutamate extruded *via* this axis could protect PCs from ferroptosis. Consistent with this hypothesis, we observed a substantial increase in extracellular glutamate levels in FOLFIRI-PCs relative to untreated cells and FOLFOX-PCs. (Fig. 3a).

**Figure 3.**
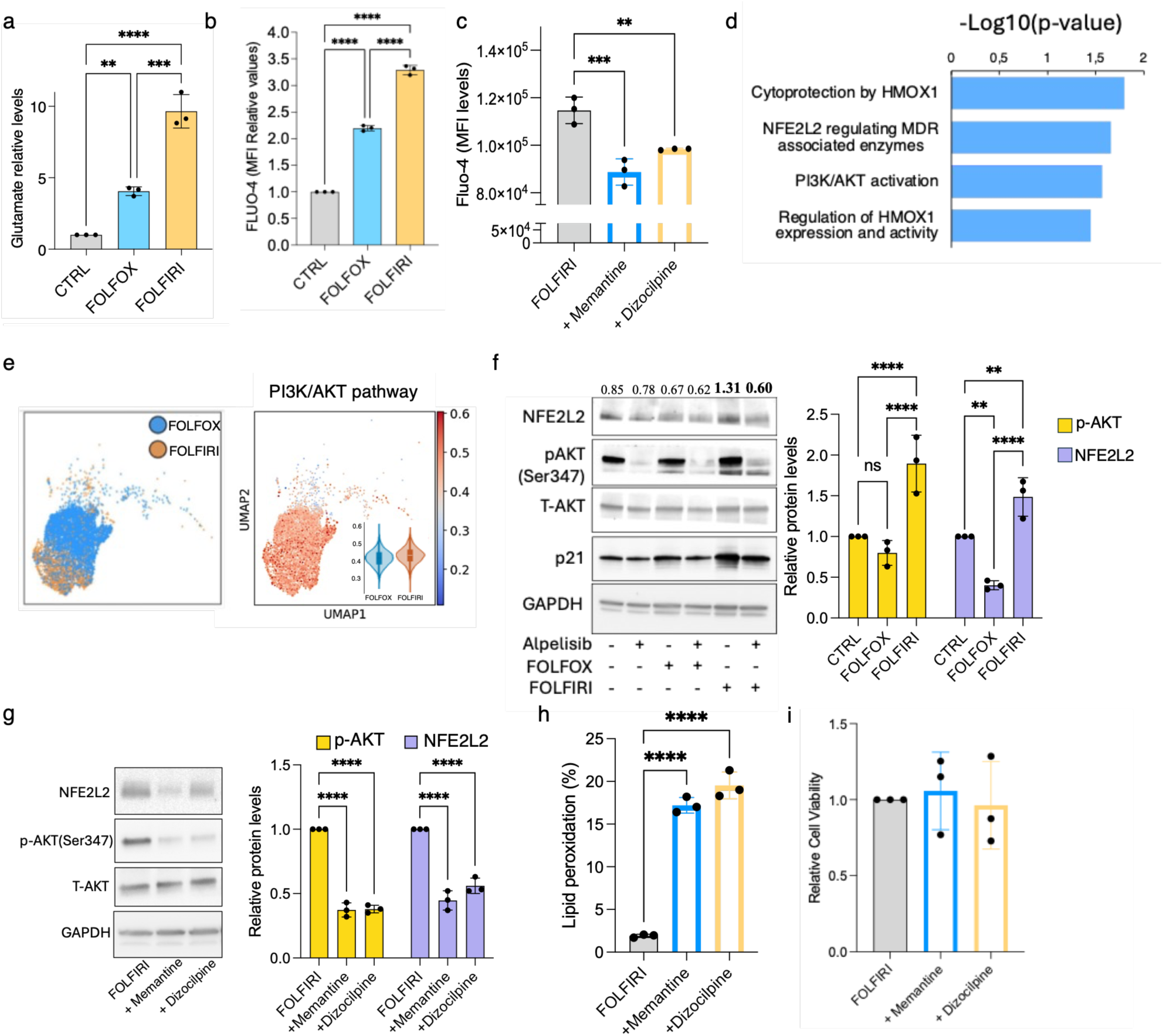
Glutamate activates NMDAR and induces cytoprotection. **a.** Glutamate secretion detected in PCs and parental CTRL. Significant differences among groups calculated by ANOVA followed by Tukey post-test (n=3 biological replicates; mean±SD; **: P<0.01; ***: P<0.001). **b.** Intracellular Ca^2+^ content quantification (expressed as MFI) in PCs and parental CTRL by flow cytometry analysis. Significant differences among groups calculated by ANOVA followed by Tukey post-test (n=3 biological replicates; mean±SD; ****: P<0.0001). **c.** Intracellular Ca^2+^ content quantification (expressed as MFI) in FOLFIRI-PCs treated with NMDAR inhibitors Memantine or Dizocilpine by flow cytometry analysis. Significant differences among groups calculated by ANOVA followed by Dunnett post-test (n=3 biological replicates; mean±SD; **: P<0.01; ***: P<0.001). **d.** Reactome analysis of genes exclusively upregulated in FOLFIRI-PCs related to PI3K/AKT and its downstream targets. **e**. UMAP embedding of sc-RNA sequencing in HCT116 PCs representing drugs and PI3K/AKT pathway for two experimental replicas. **f.** Western blot analysis and respective quantification of protein levels in parental HCT116 or PCs treated with PI3K inhibitor Alpelisib. GAPDH was used as a normalizer. Significant differences among groups calculated by ANOVA followed by Tukey post-test (n=3 biological replicates; mean±SD; **: P<0.01; ****: P<0.0001; ns: not significant). **g.** Western blot analysis of NFE2L2 and total (T-) or phosphorylated (p-) AKT in FOLFIRI-PCs treated with NMDAR inhibitors Memantine or Dizocilpine and their quantification. Significant differences among groups calculated by ANOVA followed by Dunnett post-test (n=3 biological replicates; mean±SD; ****: P<0.0001). GAPDH was used as a normalizer. **h.** Lipid peroxidation levels in FOLFIRI-PCs treated with NMDAR inhibitors Memantine or Dizocilpine. (Significant differences among groups calculated by one way ANOVA followed by Dunnett’s post-test; ****: P<0.0001). **i.** Relative cell viability of FOLFIRI-PCs treated with NMDAR inhibitors Memantine or Dizocilpine (n=3 biological replicates; mean±SD).

Glutamate is an excitatory neurotransmitter that triggers the activation of N-methyl-D-aspartate receptors (NMDARs), ushering calcium (Ca^2+^) from the extracellular milieu to the cytoplasm^31^. NMDARs are expressed not only in the central nervous system (CNS) cells, but have been reported also in several cancer types, where their role remains poorly understood^32^. In fact, based on the Human Protein Atlas (https://www.proteinatlas.org), NMDARs (GRIN1, GRIN2B, C, D, GRIN3B and GRINA) are expressed also in colon cancer and in the HCT116 CRC cells. As a proxy of the NMDAR activity, we quantified the intracellular Ca^2+^ content by flow cytometry analysis^33^, in the presence of the NMDARs inhibitors Memantine or Dizocilpine. FOLFIRI-PCs presented a higher content of Ca^2+^ than parental cells and FOLFOX-PCs (Fig. 3b). This increase was blunted by the treatment with both inhibitors (Fig. 3c), suggesting that FOLFIRI increases Ca^2+^ flux through the engagement of NMDARs.

We next asked which might be the consequences of the increase in Ca^2+^ in FOLFIRI-PCs. The concentration of Ca²⁺ in the cytoplasm is tightly regulated, and changes in Ca²⁺ concentration activate signalling events, including Protein kinase C (PKC), mitogen-activated MAPK kinases (p38 and p42/44-MAPK), and the phosphatidylinositol 3-kinase (PI3K)/AKT axis^34^. To gain insights on which pathways may be triggered by the increased intracellular Ca^2+^, we explored bulk RNA-seq data, surveying the genes specifically upregulated upon FOLFIRI treatment in PCs (Suppl. Table 1). Among the 708 genes emerging from this analysis, several downstream targets of the PI3K/AKT pathway appeared (Fig. 3d). The activation of the PI3K/AKT pathway was confirmed in our previous FOLFIRI-PCs single-cell RNA-seq data^1^ (Fig. 3e). This was further validated by western blot analysis, which revealed increased detection of both AKT phosphorylation (p-AKT) at the Ser347 residue and its downstream target, the cyclin-dependent kinase inhibitor p21, which acts as a competitor of KEAP1 for NFE2L2 binding^35^ (Fig. 3f).

Activated PI3K may prevent ferroptosis through various nodes, including mTOR, FTH1 and VDAC1^36^. However, among the pathways mostly increased upon FOLFIRI treatment, there were genes associated with the anti-ferroptotic transcription factor NFE2L2^37^ (Fig. 3d), which has been reported as a downstream target of PI3K/AKT^38^. We hence assessed whether the inhibition of PI3K/AKT may impact on NFE2L2 levels in FOLFIRI-PCs, using the specific PI3K inhibitor Alpelisib. This compound significantly reduced both AKT phosphorylation and NFE2L2 levels, thus confirming that AKT promotes the accumulation of NFE2L2 (Fig. 3f).

Our results suggest a model where, during the persistence state, glutamate binding to NMDARs increases Ca^2+^ uptake, which in turn triggers PI3K/AKT and the ferroptosis inhibitor NFE2L2^39^. To substantiate this model, we treated PCs with the NMDARs inhibitors and assessed the levels of p-AKT and NFE2L2 proteins. Indeed, both inhibitors reduced pAKT and NFE2L2 levels (Fig. 3g). Moreover, the combination of NMDARs inhibitors with FOLFIRI increased lipid peroxidation (Fig. 3h) but had no effect on cell viability (Fig. 3i). These results reinforce the notion that multiple anti-oxidative mechanisms cooperate after FOLFIRI exposure; consequently, alternative or combined approaches are necessary to trigger ferroptosis in FOLFIRI-PCs.

### NFE2L2 inhibition resensitizes PCs to ferroptosis

How does NFE2L2 drive an antioxidant effect to protect PCs from peroxidation-dependent cell death? To answer this question, we performed an RNA-seq analysis in NFE2L2 silencing conditions. Beyond the significant down regulation of known target genes, such as NQO1 (NADPH quinone dehydrogenase 1) and genes involved in pentose phosphate pathway (i.e. TKT, TALDO1 and G6PD)^40^ (Suppl. Fig. 3a), we were surprised to find the upregulation of Endoplasmic Reticulum (ER) stress and Unfolded Protein Response (UPR – PERK/ATF4) genes (Suppl. Fig. 3b).

It has been recently reported that the junction between ER and mitochondria (M), namely EM Contact (EMC), modulates lipid peroxidation and ferroptosis and that ER stress and UPR are tightly involved in EMCs regulation^41^. The close proximity between ER and M (thus, reduced distance) favours ferroptosis, while an increased space between these two compartments protects cells^41^. The molecular mechanism regulating EMC remains elusive.

We analysed transmitted electron microscopy (TEM) images in PCs and measured ER-M distance. FOLFOX treatment profoundly reduced ER-M distance, much more than FOLFIRI, as the gap separating the ER and mitochondria shortened from an average of 17 nm in untreated cells to 6 nm in response to FOLFOX and only to 12 nm in FOLFIRI-PCs (Suppl. Fig. 3c).

We wondered whether NFE2L2 may exert a role in modulating EMC. We found that NFE2L2 silencing alone halved the ER-M distance in FOLFIRI-PCs (Fig. 4a-b). Notably, Erastin by itself did not affect EMC, while the combinatorial treatment of shNFE2L2 and Erastin greatly increased EMC (Fig. 4a-b).

**Figure 4.**
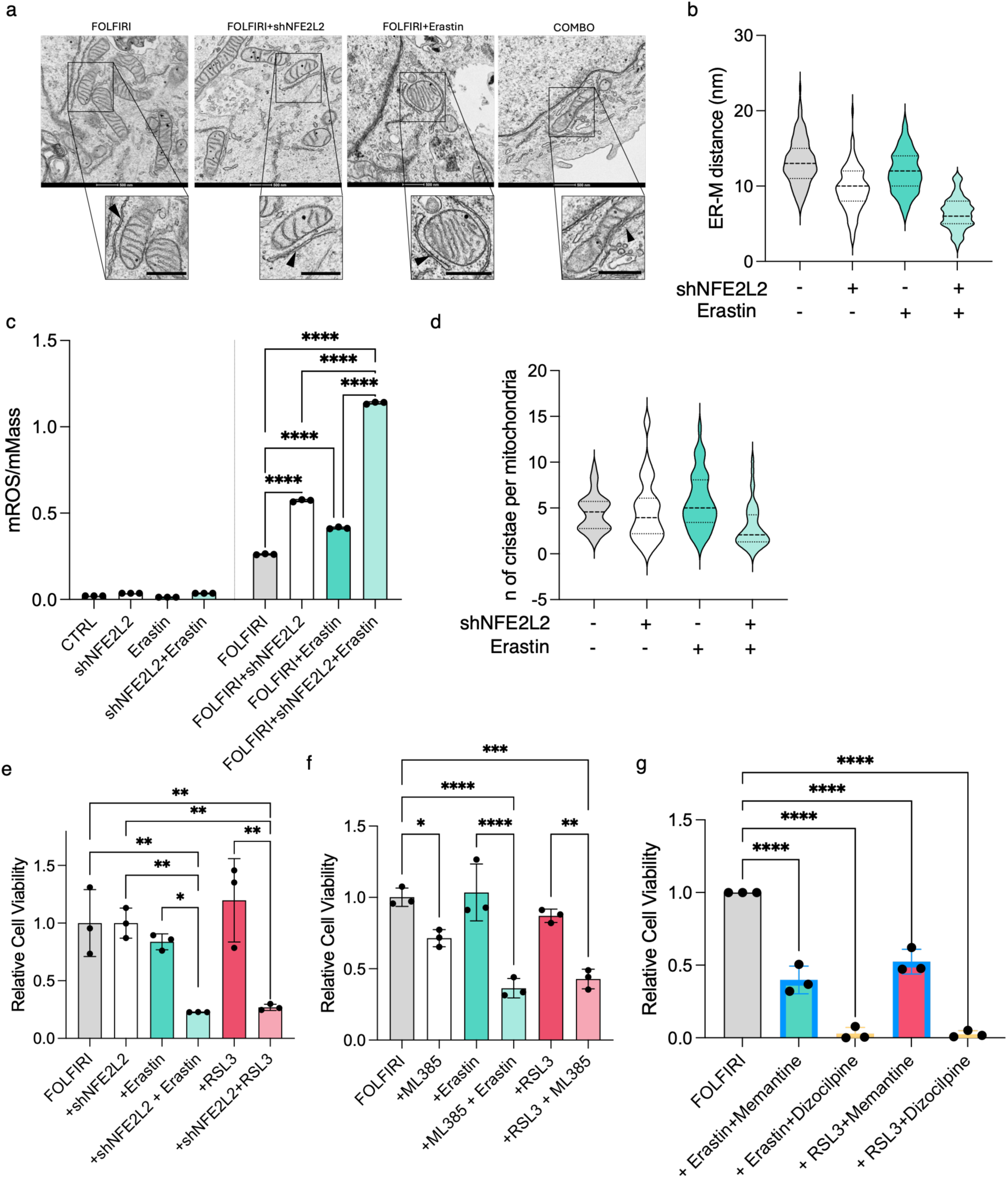
NFE2L2 regulates ER-M distance and resensitize PCs to ferroptosis. **a-b.** Representative TEM images (**a**) and corresponding quantification (**b**) of mitochondria/endoplasmic reticulum distance expressed in nm. Scale bar: 500 nm. (n=20 cells; median, first and third quartile are shown). **c**. Mitochondrial ROS quantification in CTRL cells or FOLFIRI-PCs treated with Erastin and in shNFE2L2 condition. Significant differences among groups calculated by ANOVA followed by Tukey post-test (n=3 biological replicates; mean±SD; ****: P<0.0001). **d.** Number of cristae quantification (n=20 cells; median, first and third quartile are shown). **e-f.** Relative cell viability of FOLFIRI-PCs genetically silenced for NFE2L2 (**d**) or treated with NFE2L2 inhibitor (ML385) (**e**) and treated with ferroptosis inducers Erastin and RSL3. Significant differences among groups calculated by ANOVA followed by Tukey post-test (n=3 biological replicates; mean±SD; *: P<0.05; **: P<0.01; ***: P<0.001; ****: P<0.0001). **g.** Relative cell viability of FOLFIRI-PCs treated with NMDAR inhibitors Memantine or Dizocilpine and ferroptosis inducers Erastin and RSL3. Significant differences among groups calculated by ANOVA followed by Tukey post-test (n=3 biological replicates; mean±SD; ****: P<0.0001).

The proximity between ER and mitochondria leads to the expansion of mitochondrial lipid peroxidation, inducing mitochondrial ROS (mROS) production^41^. We thus explored whether increased EMC in PCs is associated with mROS production. Consistent with the closer contact between the ER and mitochondria, the elevation of mROS was significant in FOLFOX-PCs but not in FOLFIRI-PCs (Suppl. Fig. 3d), thereby supporting the hypothesis of the lower susceptibility of FOLFIRI-PCs to mitochondrial oxidative stress. To validate the involvement of NFE2L2 in modulating this intricated cellular mechanism, based on EMC regulation and mROS expansion, we knocked down NFE2L2 in FOLFIRI-PCs and measured mROS levels, normalized to mitochondrial mass. Notably, while mROS were only moderately elevated by either shNFE2L2 or Erastin alone, the combined treatment triggered a dramatic mROS surge (Fig. 4c), likely overcoming cellular detoxifying defences and precipitating irreversible PCs damage and cell death.

Considering that the mitochondrion operates as both a major driver (through mROS generation) and a casualty of the ferroptotic cascade, we explored mitochondrial function in these cells after genetic knockdown of NFE2L2 and Erastin. During ferroptosis, mitochondria undergo distinct morphological shifts, such as condensed inner membranes, outer mitochondrial membrane blistering, and cristae depletion^42–43^. We determined that a significant loss of mitochondrial cristae was detected specifically upon the concomitant administration of Erastin and NFE2L2 knockdown (Fig. 4d), reinforcing the notion that only this combinatorial approach potently induces ferroptotic cell death.

We next explored whether the inhibition of NFE2L2 may reduce cell viability. Despite the effect of NFE2L2 knockdown on EMC, no reduction in cell viability was evident (Fig. 4e). We have shown that interfering with the Xc^-^ antiporter (using Erastin) and GPX4 (with RSL3), despite the activation of some pro-ferroptotic markers (Fig. 1a) had no impact on cell survival (Fig.1b). We next explored the possibility that targeting both paths (the Xc^-^ antiporter or GPX4 on one side, and EMC controlled by NFE2L2 on the other) may affect cell viability. The combined knockdown of NFE2L2 with either Erastin or RSL3 deeply reduced cell survival (Fig. 4e). Of note, this combined effect was not observed in FOLFOX-PCs, where the single administration of RSL3 or Erastin was sufficient to reduce viability, with no additive effect seen with NFE2L2 knockdown (Suppl. Fig. 3e). We confirmed these results also with ML385, a compound which blocks heterodimerization of NFE2L2 with small Maf proteins^44^ (Fig 4f).

To broaden the translational potential of these findings, we evaluated the combination of Erastin with NMDARs inhibitors (Memantine or Dizocilpine); as a control, we also tested the genetic silencing of GRIN1, the critical subunit of NMDARs. FOLFIRI treatment increased GRIN1 protein levels (Suppl. Fig. 3f), further supporting the model whereby cancer cells exposed to FOLFIRI treatment relies on NMDARs to prevent ferroptosis and survive. As seen with NMDARs inhibitors, also GRIN1 silencing significantly reduced NFE2L2 levels (Suppl. Fig. 3f). Importantly, while neither NMDAR inhibition (Fig. 3i) nor GRIN1 genetic silencing (Suppl. Fig. 3g) alone was sufficient to reduce the viability of PCs, the combination with Erastin or RSL3, triggered cell death in FOLFIRI-PCs (Fig. 4g). This effect was observed exclusively following persistence, and not in the parental cells (Suppl. Fig. 3h), sustaining the hypothesis that this phenotype is specifically induced by FOLFIRI treatment.

### Mitochondrial ROS precipitate ferroptosis induced by Xc^-^ antiporter inhibition

Our findings demonstrate that FOLFIRI-PCs trigger two anti-ferroptotic programs. One pathway depends on the recruitment of the cystine-glutamate antiporter Xc^-^ which activates the GSH/GPX4 cascade. However, targeting this system alone fails to sensitize cells to ferroptosis because of the parallel NMDARs/NFE2L2/mitochondria axis. Through NFE2L2 regulation, FOLFIRI-PCs restrict mitochondrial impairment and control oxidative stress to support survival.

Iron is a vital catalyst in this process. Free intracellular iron both reacts with hydrogen peroxides *via* the Fenton reaction to generate highly reactive hydroxyl radicals, which rapidly accelerate the oxidation of lipids^45^ and plays a critical role serving as a fundamental component in the biosynthesis of heme and iron−sulfur (Fe−S) clusters, which serve as crucial cofactors within the mitochondrial respiratory chain^46^. Nonetheless, a striking accumulation of Fe^2+^ was observed in FOLFIRI-PCs and this increase promoted survival rather than cell death (Fig. 1d). We hypothesized that in FOLFIRI-PCs, most of the Fe^2+^ is directed toward mitochondria to sustain organelles fitness, essential for cell survival. To test this hypothesis, we first evaluated mitochondrial Fe^2+^ levels using Mito-FerroGreen, a fluorescent probe that detects Fe^2+^, co-stained with MitoBright, a mitochondria-retained dye. We confirmed that FOLFIRI-PCs exhibited a marked increase in mitochondrial Fe^2+^ compared to both control and FOLFOX-PCs (Fig. 5a-b) which is prevented by DFO, that severely impaired PC survival (Fig. 1d). Moreover, Rotenone (RTN), a potent mitochondrial complex I inhibitor, significantly decreased FOLFIRI-PC viability and elevated lipid peroxidation, consistent with the effects due to DFO (Fig. 5c-d). These findings suggest that the increase in Fe^2+^ detected in FOLFIRI-PCs is mostly directed to the mitochondria machinery, thus preserving its function and protecting PCs from peroxidation of the lipid membranes.

**Figure 5.**
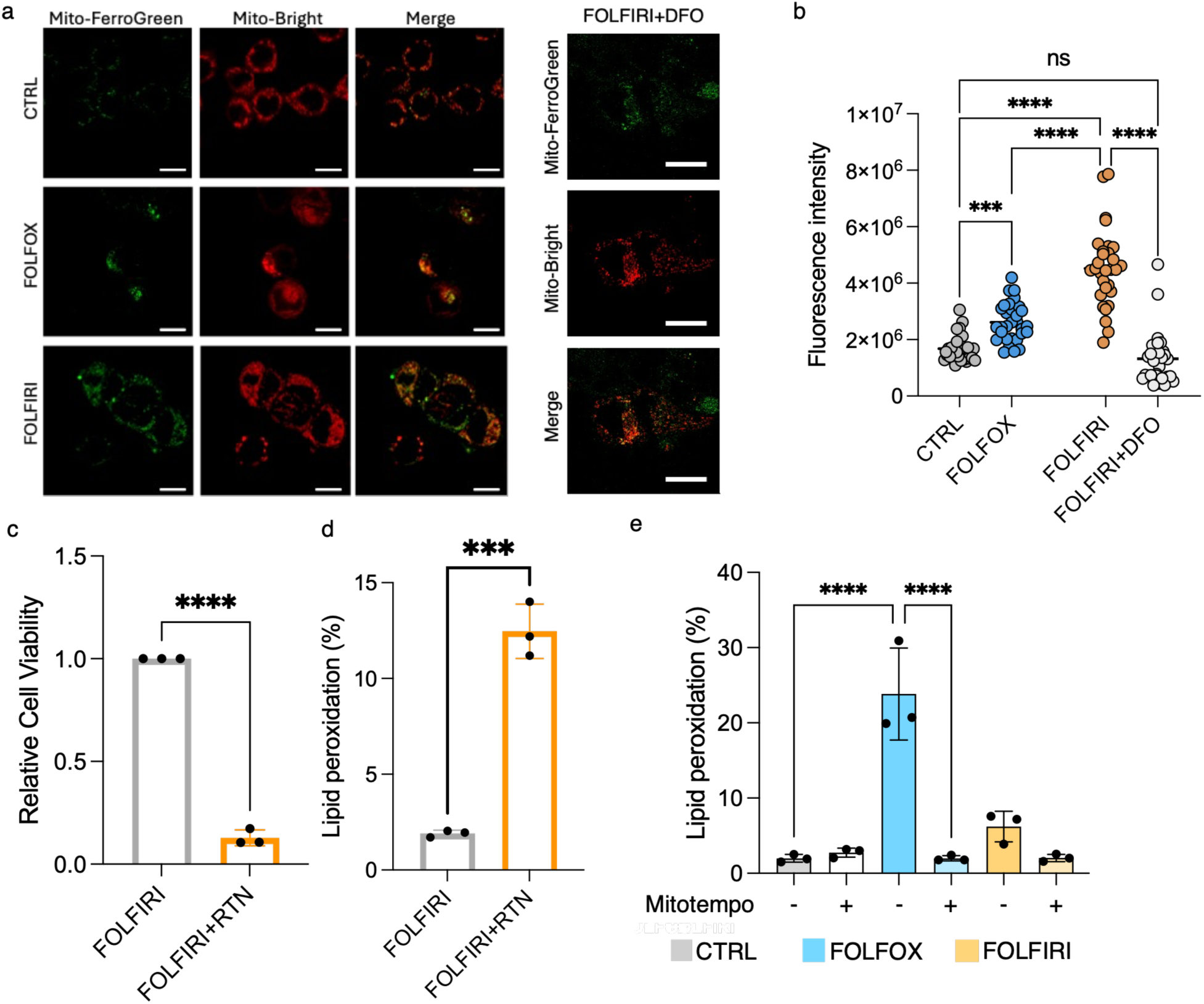
Mitochondrial ROS precipitate ferroptosis induced by Xc^-^ antiporter inhibition. **a-b.** Representative images and fluorescent microscopy analysis at confocal resolution of CTRL, FOLFOX- or FOLFIRI-and FOLFIRI+DFO-PCs (**a**) for the detection of Fe^2+^ (Mito-FerroGreen) or mitochondrial compartment (Mito-Bright). Significant differences among groups (**b**) calculated by ANOVA followed by Tukey post-test (n=3 biological replicates biological replicas and 10 cells per condition analysed; mean±SD; ***: P<0.001; ****: P<0.0001). **c-d.** Relative cell viability (**c**) and lipid peroxidation quantification (**d**) of FOLFIRI-PCs treated with Rotenone. Significant differences among groups calculated by Student’s t-test (n=3 biological replicates; mean±SD; ***: P<0.001; ****: P<0.0001). **e.** Lipid peroxidation levels quantified as percentage positivity (%) in PCs or parental CTRL treated with Mitotempo. Significant differences among groups calculated by ANOVA followed by Tukey post-test (n=3 biological replicates; mean±SD; ****: P<0.0001).

Conversely, the restriction of mitochondrial Fe^2+^ (Fig. 5a-b), alongside Xc^-^ activity blockade and impaired NADPH synthesis (Fig. 2b-c), corroborates the idea that Fe^2+^ is mostly funnelled into the pro-ferroptotic oxidative pathway within FOLFOX-PCs. To confirm the central role of the mROS in the ferroptosis induction, FOLFOX-PCs were treated with the specific mROS scavenger Mitotempo. This treatment mitigated both lipid peroxidation and mROS levels, thereby rescuing cell viability during ferroptotic challenge (Suppl. Fig. 5b-d). Overall, these findings underscore the central role of mitochondrial ROS generation in ferroptotic sensitization. During FOLFIRI exposure, NFE2L2 acts as a detoxification gatekeeper by maintaining mitochondrial distance and mitigating mROS production. In turn, accumulated mROS drive membrane lipid peroxidation, counteract the Xc^-^ antioxidant defence, and precipitate ferroptosis. Consequently, the simultaneous inhibition of both the Xc^-^ system and NFE2L2-mediated mitochondrial regulation is strictly required to restore ferroptotic sensitivity and trigger irreversible cellular destruction in FOLFIRI-PCs.

### Sulfasalazine in combination with NFE2L2 inhibition ablates FOLFIRI-PCs

Although Erastin and RSL3 have been used to study the induction of ferroptosis *in vitro* and *in vivo*, they never reached the clinics, due to toxic effects recorded in preclinical studies^47–48^.

We then evaluated Sulfasalazine, an anti-inflammatory and immunomodulatory agent approved for chronic inflammatory diseases^49^ and recently investigated in clinical trials for metastatic colorectal cancer, glioblastoma, and acute leukaemia. Furthermore, Sulfasalazine has been repurposed as an inhibitor of the Xc^-^ antiporter system to trigger ferroptosis^50–51^. Sulfasalazine, at a dose reducing survival by 30% of HCT116 parental cell, effectively increased lipid peroxidation (Suppl. Fig. 5a). We then tested the effect of sulfasalazine in the context of CRC persistence. In line with the results obtained with Erastin and RSL3, Sulfasalazine was effective in ablating FOLFOX-PCs (Fig. 6a), while having no effect in FOLFIRI-PCs (Fig. 6b). The combination of sulfasalazine with the specific NFE2L2 inhibitor ML385 resensitized FOLFIRI-PCs and strongly reduced cell viability (Fig. 6c).

**Figure 6.**
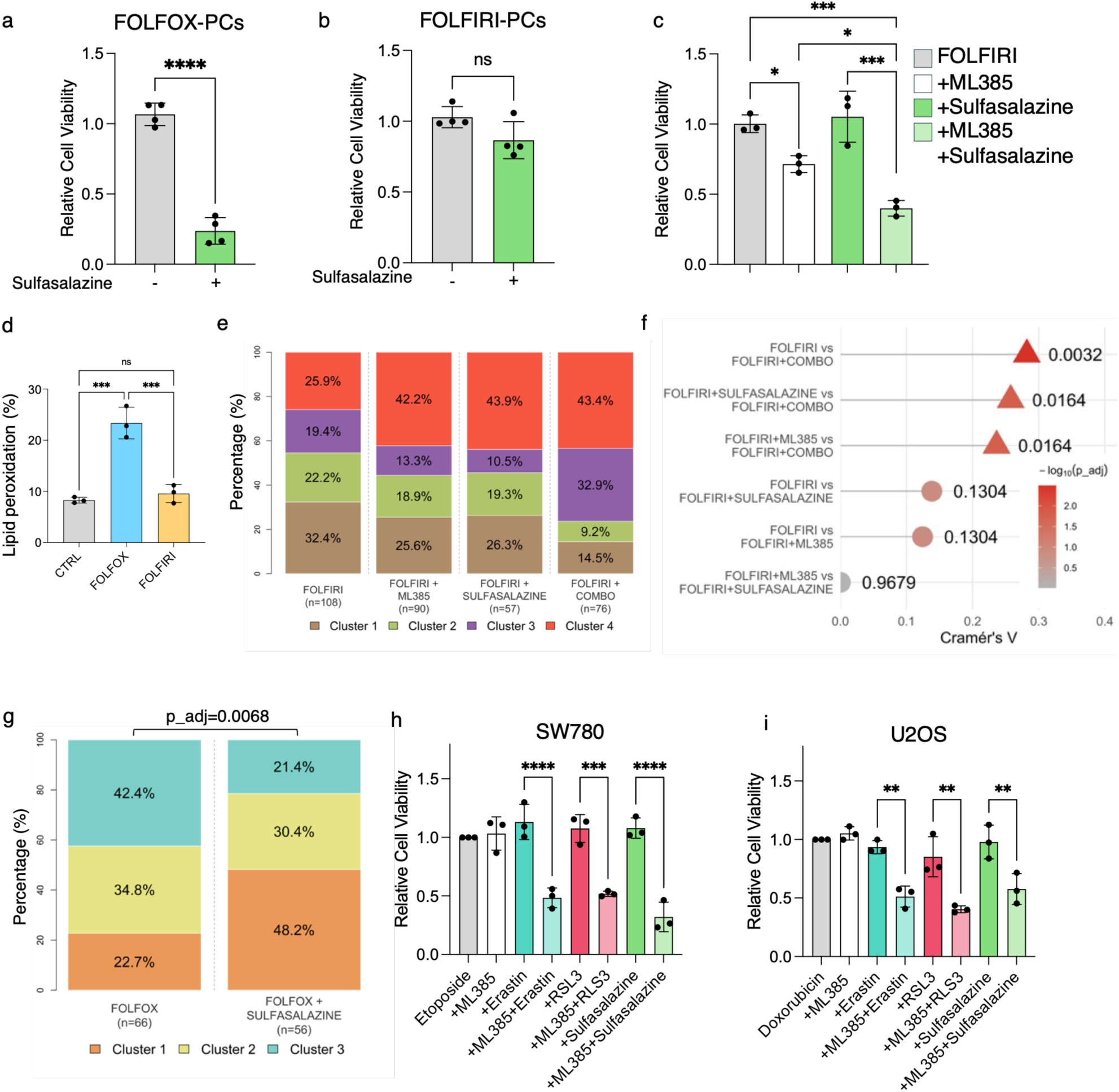
Sulfasalazine recapitulates ferroptosis induction and inhibits PCs. **a-b.** Relative cell viability of FOLFOX-(**a**) or FOLFIRI-PCs (**b**) upon Sulfasalazine treatment. Significant differences among groups calculated by two-tailed Student’s t-test (n=4 for b biological replicates; mean±SD; ****: P<0.0001; ns: not significant). **c.** Relative cell viability of FOLFIRI-PCs treated with NFE2L2 inhibitor ML385 and sulfasalazine. Significant differences among groups calculated by ANOVA followed by Tukey post-test (n=3 biological replicates; mean±SD; *: P<0.05; ***: P<0.001). **d.** Lipid peroxidation levels quantified as percentage positivity (%) in parental patient-derived organoids (PDOs) CTRL or PCs obtained by chemotherapy exposure by flow cytometry analysis. Significant differences among groups calculated by ANOVA followed by Tukey post-test (n=3 biological replicates; mean±SD; ***: P<0.001; ns: not significant). **e-g.** Abundance of phenotypic clusters and respective significance obtained by PDOs segmentation upon FOLFIRI-based (e-f) or FOLFOX-based treatments quantified and visualized *via* a stacked barplot. Adjusted BH *p*-values and Cramer’s V effect size are reported. **h-i.** Relative cell viability of SW780 (**h**) and U2OS (**i**) PCs treated with etoposide or doxorubicin, respectively, ferroptosis inducers and ML385. Significant differences among groups calculated by ANOVA followed by Tukey post-test (n=3 biological replicates; mean±SD; ***: P<0.001; ****: P<0.0001).

To extend these observations to an experimental setting closer to the clinic, we used a CRC patient-derived organoid (PDO) derived from a patient affected by metastatic CRC to the liver, exposed for two weeks to either FOLFOX or FOLFIRI to induce persistence. Also, in this PDO, FOLFOX, but not FOLFIRI, increased lipid peroxidation in PCs (Fig. 6d), with the combination of FOLFOX with Sulfasalazine reducing PDOs area (Suppl. Fig. 5b-c). Conversely, the combination of Sulfasalazine (as Xc^-^ antiporter system inhibitor) and ML385 (NFE2L2 inhibitor), was required to significantly affect FOLFIRI-PCs (Suppl. Fig. 5b-d). To confirm the effect on cell death, we applied a deep-learning pipeline extracting representative phenotypic states from PDOs^52^. Unsupervised clustering analysis revealed four clusters in FOLFIRI- and three clusters in FOLFOX-based therapies (Suppl. Fig. 5e-f). Ferroptosis-based treatments in FOLFIRI-PCs altered clusters abundance, leading to a widespread reduction in Clusters 1 and 2 (highly structured and regular organoid architectures) and an increase in Cluster 4 (intermediate phenotype). Notably, Cluster 3, the most extreme degenerate and ’sprouting’, enriched significantly only following doublet pro-ferroptosis therapies (Fig. 6e-f). Similar results were obtained in the FOLFOX-based treatments, with a prevalence of the deranged cluster phenotypes in the combination (Fig. 6g). Altogether, these findings demonstrate a significant disruption of the organoids architectural structure, thus supporting a potent cell death phenotype following combinatorial treatments against both Xc^-^ antiporter and NFE2L2.

Topoisomerase inhibitors are widely used in oncology, for the treatment of colorectal^53^, pancreatic^54^, ovarian^55^, osteosarcoma^56^ and bladder cancer^57^, among the others, as first line therapy for metastatic disease or in case of resistance to previous regimens. We then assessed whether other topoisomerase inhibitors displayed the same phenotype seen with SN38, in cell lines derived from tumour types where these compounds are routinely used, including the SW780 bladder cancer cell line and the U2OS osteosarcoma model using Etoposide and Doxorubicin, respectively. Both cell lines were treated with Erastin, RSL3 and Sulfasalazine at dosages able to reduce viability in the parental counterpart (Suppl. Fig. 5g-h). None of the combinations was able to reduce cell viability on itself in PCs (Suppl. Fig. 5i-j) while the combination with NFE2L2 inhibitor ML385 was able to reduce viability (Fig. 6h-i), as showed for the CRC cancer models. These results suggest that resistance to ferroptosis induction is specific of topoisomerases inhibition and provides guidance to ameliorate patients’ response to standard treatments.

## Discussion

Tumour cell death is the main goal of most anti-cancer therapies^58^, as well as eradication of tumour reservoirs, such as persister cells^59^. In this study, we show that ferroptosis is strongly activated during chemotherapy-induced persistence of colorectal cancer cells, with platinum-based combination sensitizing persister cells to ferroptosis induction. Surprisingly, following topoisomerase inhibition, persister cells were instead resistant to the induction of ferroptosis. In these cells, CD44 cooperates with SLC7A11, the main subunit of the Xc^-^ antiporter system, increasing cystine uptake and activating the GPX4 antioxidant pathway. Notably, blocking the Xc^-^ system, by Erastin or the GPX4 inhibitor RSL3, was not sufficient to sensitize cells to ferroptosis, suggesting the involvement of an additional mechanism triggering resistance. We found an increase of the extracellular glutamate which, in turn, stimulates the NMDARs to activate the downstream NFE2L2 signalling axis which, by directly maintaining ER-mitochondrial distance, mitigates oxidative stress. Ultimately, targeting both axes simultaneously is essential to overcome this defence mechanism, induce ferroptosis, and clear the persister cell population.

The role of the synaptic plasticity receptors NMDARs in cancer is starting to emerge. NMDARs exert a central role in the synaptic transmission and plasticity^60^. However, recent evidence suggests that these receptors are expressed in tissues of different tumour types, such as pancreatic^61–62^, small cell lung^63^ and breast cancer^64^. Aberrant accumulation of glutamate in the extracellular compartment activates glutamate receptors on cancer cells, promoting malignant growth^65^. NMDARs stimulate several tumorigenic pathways, such as mTOR, ERK1/2, and CREB through Ca^2+^ uptake^32^. We propose that NMDARs, when activated by glutamate binding, mediate ferroptosis resistance in response to topoisomerase inhibitor treatments. Our finding is consistent with existing data that correlate these classical synaptic receptors with rituximab resistance in neurological disorders such as autoimmune encephalitis or Ca^2+^-dependent excitotoxicity in epilepsy^66–67^. In addition, NMDARs have been associated with refractory to temozolomide chemotherapy in glioblastoma patients, due to the increase of the DNA repair enzyme, O6-methylguanine DNA methyltransferase^68^. Our data demonstrate that NMDARs, by activating the Ca^2+^-dependent survival signals, such as AKT phosphorylation, trigger the NFE2L2 transcription factor in the context of cancer persistence.

Endoplasmic reticulum-mitochondria contact (EMC) is emerging as the central hub in mitochondrial ROS production and for the oxidative ferroptotic hotspot^41^. Several proteins, by acting as membrane tethers, have been described to maintain EMC^69^. However, the driving factors behind this regulation remain poorly defined. Our finding sheds light on the role of NFE2L2 in triggering this process. We hypothesize that NFE2L2 controls ER stress and the unfolded protein response and, by maintaining the distance among organelles, halts both lipid peroxidation and mitochondrial damage. Such a complex regulation confers resistance to ferroptosis and opens new avenues for the characterization of the mechanisms governing EMC, also beyond the setting of persistence.

Targeting iron metabolism, either through Fe^2+^ deprivation or by leveraging Fe^2+^ accumulation to trigger ferroptosis, represents a promising therapeutic strategy in oncology^70^. As such, increasing cellular Fe^2+^ uptake is a well-established strategy to induce ferroptosis, also in persister cells^30,71^. Conversely, the potential protective role of Fe^2+^ in PCs has not been previously investigated to the best of our knowledge. Indeed, despite its role in driving ferroptosis, Fe^2+^ could alternatively promote cancer cell growth, a phenomenon already documented in signalling pathways associated with other forms of programmed cell death^72–73^. Here, we demonstrate that following FOLFIRI treatment, Fe^2+^ chelation ablates PCs. In several settings Fe^2+^ chelation has been proposed as a strategy to treat cancers, for example in colorectal, ovarian and hepatocellular carcinoma, glioblastoma and leukemia^70^. We herein propose that Fe^2+^ chelation may be exploited to interfere with the survival of PCs arising from topoisomerase treatments.

Why do these PCs require iron? We argue that Fe^2+^ in this setting is required for mitochondrial fitness. In particular, we delineate the role of Fe^2+^ in maintaining mitochondrial homeostasis, which is critical for tumour initiation and growth^74^. Indeed, beyond the classical role in the Fenton reaction, which generates highly reactive hydroxyl radicals and drives lipid peroxidation^75^, Fe^2+^ can be exploited by cancer cells to sustain mitochondrial Fe-S cluster and heme synthesis^76^. In this study, we propose that this distinctive mitochondrial mechanism acts as an essential player in sustaining cell survival during treatment and cancer persistence. To support this hypothesis, we speculate that NFE2L2, by modulating the labile Fe^2+^ pool via FTH1 regulation, ensures an adequate iron supply while preventing excess Fe^2+^ from initiating detrimental Fenton reactions^77^. Conversely, an inability of cancer cells to compartmentalize Fe^2+^ triggers a massive surge in mitochondrial reactive oxygen species. Consequently, employing an inhibitor of Fe^2+^ transport to mitochondria, or utilizing a combination of drugs that deprives the cancer cell of mitochondria protective mechanisms, forces it into rapid, catastrophic, iron-dependent cell death.

Exploiting the mechanistic links between chemotherapy and iron-dependent pathways during the persistent state provides a promising framework for developing synergistic combination regimens. Accordingly, we suggest that ferroptotic induction could represent a successful strategy in combination with FOLFOX. Conversely, exploiting the ferroptotic pathway after topoisomerase inhibitor treatments is more challenging, as it requires the simultaneous inhibition of both antioxidant strategies, namely the Xc^-^ antiporter system and the NMDAR/NFE2L2 axis, to dismantle the antioxidant response and eradicate FOLFIRI persister cells.

How to safely induce ferroptosis for cancer treatment remains a critical question, as the primary challenge lies in developing a therapeutic strategy that protects healthy cells while selectively targeting tumour cells, alongside identifying patients who would most benefit from ferroptosis-based interventions^78^. To address this, we successfully utilized Sulfasalazine, an FDA-approved anti- inflammatory drug, which possesses well-characterized pro-ferroptotic properties^51^. Although several studies have already evaluated its efficacy as an antitumor agent, including in colorectal cancer^79^, our data repurpose its application to the context of cancer persistence. Specifically, we combine it with a targeted therapy against NFE2L2 to overcome drug resistance, validating these findings in a highly clinically relevant model, such as colorectal cancer patient-derived organoids that recapitulate the FOLFIRI-induced activation of the antioxidant response.

In conclusion, we propose exploiting iron-dependent mechanisms to eradicate persister cells in colorectal cancer, while also accounting for the diversity of the plastic response to therapy as an intrinsic property of these cells. Alternative approaches must be tailored to successfully target survival machineries during the persistent state. Furthermore, additional analyses are required to develop patient stratification strategies, identifying individuals who would benefit most from ferroptosis induction versus those who could alternatively receive combination therapies designed to disrupt the redox balance and halt tumour progression.

## Methods

### Cell line and Patient derived organoids

Experiments were performed in CRC HCT116, SW780 AND U2OS cell lines, maintained in media composed by DMEM, 10% FBS and 1% Penicillin-Streptomycin, as recommended by supplier. Cell line authentication was performed by Thermo Fisher, AmpFlSTR® Identifiler® Plus PCR Amplification Kit. All cell lines were tested for mycoplasma and resulted negative.

Samples from liver metastatic gastrointestinal cancers were obtained from San Raffaele Hospital (Milan, Italy). Patient Derived Organoids (PDO) cultures were established as previously reported^28^. Briefly, fresh tissues were minced immediately after surgery, conditioned in PBS with 5 mM EDTA, and digested for 1 hour at 37°C in a solution containing 2× TrypLE Select Enzyme (ThermoFisher) in PBS with 1 mM EDTA and DNAse I (Merck). Cell release was facilitated by pipetting. The dissociated cells were then collected, suspended in 120 μl of growth factor-reduced Matrigel (Corning 356231, Fisher Scientific), and seeded in single domes in a 24-well flat-bottom cell culture plate (Corning). After the domes solidified, they were covered with 1 ml of complete human organoid medium. The medium was replaced every 2–3 days.

### Persister cells derivation and treatments

To obtain persister cells, 1.2×10^6^ HCT116 cells were plated on a 6 cm plate and continuously exposed for two weeks to FOLFOX: 5-FU 10 µM and oxaliplatin 1 µM or FOLFIRI: 5-FU 10 µM and SN38 10 nM. Medium and drugs were refreshed every four days. HCT116 Persister cells were treated for 72 hours with Erastin (SelleckChem S7242), RSL3 (SelleckChem S8155) or Sulfasalazine (SelleckChem S1576) at respective concentrations of 5 µM, 2.5 µM and 500 µM. SW780 cells were treated once week with 10 µM Etoposide (SelleckChem S1225); SW780 persister cells were treated for 72 hours with 500 nM Erastin, 25 nM RSL3 or 50 µM Sulfasalazine. Doses for treatments were chosen based on the half-maximal inhibitory concentration (IC50) at 72h. U2OS cells were treated twice a week with 10 nM Doxorubicin (SelleckChem E2516). U2OS persister cells were treated for 72 hours with 500 nM Erastin, 250 nM RSL3 or 50 µM Sulfasalazine. Chemonaive cells were used as a control group. ML385 (SelleckChem S8790) was used at a final concentration of 1 µM for 72 hours. Persister cells maintained in culture for 72 hours with complete medium were used as control. Mitotempo (MedChemExpress HY-112879) or Rotenone (R8875; Merck) were used for 48h at 10 µM and 1 µM concentrations, respectively. Deferoxamine (MedChemExpress HY-B1625) was used for 48h hours at 1 µM concentration. Alpelisib (MedChemExpress HY-15244) was used at 5 µM concentration for 16h. Memantine (MedChemExpress HY-B0591) and Dizocilpine (MedChemExpress HY-15084B) were used at 10 µM for 72h.

### PDOs drug treatment and segmentations

PDOs were treated for two weeks with FOLFOX: 10 µM 5-FU and 10 µM of oxaliplatin or FOLFIRI: 10 µM 5-FU and 10 nM SN38. Medium and drugs were refreshed every four days. After induction of persistence, PDOs were treated with sulfasalazine (500 µM) and ML385 (1 µM). Images of the PDOs were acquired at the inverted microscope and were analysed using ImageJ. Briefly, three to five fields per well were acquired and the area of each organoid was measured. In addition, 74 images (1.02 μm/px, 8-bit, 1338 x 1040px-sized images, acquired through AxioCamMR3) encompassing FOLFIRI- and FOLFOX-based treated PDOs were processed through Napari Organoid Analyzer, a deep learning pretrained model consisting of Yolov3+ and SAM network for the automatic segmentation of single PDO instances in the image^52^. An expert-led quality check was performed on automated segmentation masks. According to the therapy delivered, the number of PDO instances segmented resulted in: FOLFIRI (n=101, 14 images), FOLFIRI+ML385 (n=80, 15 images), FOLFIRI+SULFASALAZINE (n=51, 11 images), FOLFIRI+COMBO (n=63, 14 images), FOLFOX (n=54, 13 images), and FOLFOX+SULFASALAZINE (n=46, 7 images). For each PDO instance, automatic feature extraction was performed through convolutional deep neural networks, namely ConvNeXt-tiny and DenseNet-121^80–81^. Subsequently, an Autoencoder was trained to compress feature space while minimizing reconstruction loss. On the generated latent space, the number of clusters was selected among k = {3,4,5} through Consensus Clustering implemented in Bioconductor ConsensusClusterPlusR package, by using Gaussian Mixture Model and Partitioning Around Medoids (PAM) as independent estimators. Upon establishing the optimal *k*, the PAM algorithm was deployed to identify the *k*-medoids and assign definitive cluster labels to all individual PDO instances across the entire dataset. A global contingency table (*clusters* × *t*ℎ*erapies*) was constructed to map the occurrence of phenotypic clusters across all therapeutic conditions, and global statistical association was assessed using the Chi-squared (*χ*2) test. To dissect specific treatment effects, therapies were further compared via pairwise *χ*2 analyses using reduced (*clusters*×2) contingency frameworks by reporting *p*-values and Cramer’s V effect size.

### Cell viability assay

Cell viability was assessed using crystal violet staining. Briefly, after removing the supernatant, the cells were washed with PBS. Crystal violet was then added to live cells and incubated for 15 minutes. Images were captured using an inverted microscope and analysed with ImageJ software to quantify cell viability.

### Labile Iron quantification

To detect the Labile Iron Pool (LIP) 1,5×10^5^ PCs were plated in 6 wells and then incubated with PBS and Calcein AM (acetoxymethyl ester) 50 nM at 37°C for 30’ in the dark. Mean fluorescence intensity (MFI) of 20’000 acquired events was detected with CytoFLEX S (Beckman Coulter) and analysed with FlowJo (Becton Coulter) (Ex/Em:495/521 nm). The MFIs were then normalized to the unstained samples to account for autofluorescence. LIP was calculated as -log (Calcein Fluorescence Intensity) related to control. FerroOrange (Cell Signaling #36104) was also used for labile Fe^2+^ quantification as reported by manufactures. Briefly, cells were incubated with 1 µmol/L FerroOrange working solution for 30 min. Then, cells were harvested and 20’000 acquired events were detected with CytoFLEX S (Beckman Coulter) and analysed with FlowJo (Becton Coulter) (Ex/Em: 543 /580 nm)

### Mitochondrial ROS, Mitochondrial Mass and Mitochondrial Fe^2+^

HCT116 cells were treated for two weeks with FOLFOX and FOLFIRI as previously specified. Then, cells were incubated with MitoSOX™ Red (Invitrogen M36008; 5 μM) resuspended in HBSS for 10 min for mitochondrial ROS detection or MitoTracker Green (Invitrogen M7514; 100nM) for 20 min at 37° C in the dark. MFI of 20’000 acquired events was detected at CytoFLEX LX (Beckman Coulter) and analysed by FlowJo (Becton Dickinson v10.8.1) (MitoSOX™ Red at Ex/Em: 510/580 nm). For mitochondrial Fe^2+^ detection cell were incubated with Mito-FerroGreen (5 μmol/l; M489 Dojindo) or MitoBright (200 nmol/l; MT12-12 Dojindo) for 30 minutes at 37 °C. The cells were observed by confocal fluorescence microscopy (TCS SP5 Laser Scanning Confocal Microscope Leica × 63 magnification).

### Lipid peroxidation

BODIPY™ 581/591 C11 (Invitrogen) was used to detect lipid peroxidation in persister cells. PBS containing BODIPY at a final concentration of 2 µM was substituted to the cell culture medium, and the cells were incubated in the dark for 15 minutes at 37°C. MFI of 20’000 acquired events was detected using the CytoFLEX S (Beckman Coulter) and analysed with FlowJo (Becton Dickinson v10.8.1). In the reduced state, the excitation and emission maxima of BODIPY 581/591 C11 are 581/591 nm; after oxidation, the probe shifts the excitation and emission to 488/510 nm. Lipid peroxidation was calculated as the ratio of oxidized to non-oxidized form.

### Glutathione, NADPH, glutamate and cystine measurement

NADP+/NADPH-Glo and GSH-Glo kits (Promega) were used to measure intracellular levels of NADPH and GSH levels as reported by the manufacturer after two weeks of treatment. Glutamate secretion was quantified in the medium harvested after two weeks of treatment by using Glutamine/Glutamate-Glo (Promega). Cystine Uptake Assay Kit (UP05 Dojindo) was used for cystine quantification. Luminescence and fluorescence levels were recorded using a plate-reading luminometer (Mithras LB940) as directed by the manufacturer and normalized on cells number.

### Ca^2+^ quantification

Intracellular calcium uptake was performed by incubating 2×10^5^ persister cells with 1 μM Fluo-4 (MedChemExpress HY-101896) diluted in HHBS (without Ca and Mg) containing 0.04% Poloxamer (MedChemExpress HY-D1005) for 1hour at 37 °C. Then, cells were acquired by flow cytometry and MFI of 20’000 acquired events was detected at CytoFLEX LX (Beckman Coulter) and analysed by FlowJo (Becton Dickinson v10.8.1) (Ex/Em: 485/526 nm).

### Transmitted Electron Microscopy (TEM) assay

For TEM analysis, PCs cells were treated and acquired as previously reported^1^. Then, for measurement of ER/mitochondria distance, images acquired at Ceta CCD camera (FEI, Thermo Fisher Scientific) were analysed using ImageJ software.

### Western Blot

Samples were collected as pellets and then lysed using a lysis buffer (1M Tris-HCl, pH 6.8, and 10% SDS in H₂O) at 95°C for 10 minutes. Afterwards, the samples were sonicated (12% amplification) for 12 seconds and ultracentrifuged for 10 minutes at 13’000 rpm. Membranes were probed with following antibodies: GAPDH (GT239; GeneTex), TFRC (D7G9X #13113 Cell Signaling), FTH1 (#3998 Cell Signaling); GPX4 (#52455 Cell Signaling); NFE2L2 (E3J1V; #33649 Cell Signaling); AKT (pan; 11E7; #4685 Cell Signaling); Phospho-AKT(Ser473)(D9E XP #4060 Cell Signaling); p21 Waf/Cip1 (12D1; #2947 Cell Signaling); CD44 (E7K2Y; #37259 Cell Signaling); SLC7A11(A7C6-R; MA5-44922 Invitrogen); GRIN1 (D65B7; #5704 Cell Signaling). The membranes were then incubated with an appropriate secondary antibody for 1 hour. Signals were detected by using ChemiDoc Imaging Systems after 5 min of incubation with ECL™ Prime Western Blotting Detection Reagent (Sigma-Aldrich).

### Immunofluorescence

HT116 PCs cells were plated on slides as follows: 300’000 cells were plated on slides previously coated with Cell-Tak (Merck CLS354240) dissolved in 0.1 M NaHCO_3_. Then, cells were fixed with 4% paraformaldehyde for 10 min, permeabilized with 0.5% Triton-X, and blocked for 1 h with 5% bovine serum albumin. The primary antibodies against CD44 (1M7.8.1; GTX15834 GeneTex), SLC7A11 (A7C6-R; MA5-44922 Invitrogen) were incubated overnight at 4 °C and then for 1 h with the secondary antibody at room temperature. Slides were counterstained with 4′,6-diamidino-2-phenylindole (DAPI) for nuclei labelling and mounted on glass slides with ProLong Gold antifade reagent. Images were collected by TCS SP5 Laser Scanning Confocal Microscope (Leica) × 63 magnification.

### Proximity Ligation Assay (PLA)

300’000 cells as control or treated with FOLFIRI were infected to silence CD44 or a SCR control and then plated on slides previously coated with Cell-Tak as above. After fixing and permeabilization, cells were incubated with the primary antibodies against CD44 (E7K2Y; #37259 Cell Signaling) and SLC7A11(A7C6-R; MA5-44922 Invitrogen) overnight at 4 °C. After incubation with the probes and ligation and amplification, slides were mounted by using in situ mounting media with DAPI and acquired at 100x magnification by Axio Imager A2 fluorescence microscope (Zeiss). ImageJ was used to process images. The PLA puncta were isolated from the background and then counted. The results were expressed as PLA puncta/nuclei. Significant differences among groups were calculated by applying ANOVA followed by Tukey post-test.

### Apoptosis evaluation

HCT116 PCs were harvested and stained with ANNEXIN V-FITC (Invitrogen - eBioscience Annexin V-FITC Apop kit; BMS500FI) as recommended by the supplier. Cells were acquired at CytoFLEX LX (Beckman Coulter) and analysed by FlowJo (Becton Dickinson v10.8.1).

### RNA interference

For genetic silencing, HCT116 cells were infected with specific inducible small hairpin RNA (shRNA) targeting NFE2L2 (Addgene plasmid # 136584; http://n2t.net/addgene:136584; RRID: Addgene136584) cloned into PLK0 vector. As a control we used shRNA targeting luciferase, shLuc (Addgene plasmid # 136587; http://n2t.net/addgene:136587; RRID: Addgene136587) cloned into PLK0 vector. Lentiviral particles were added to HCT116 cells, together with 4 μg/mL polybrene (Sigma) for 16 hours. After 48 hours medium was replaced and 2μg/mL of puromycin was added and maintained along chemotherapy treatments. After the cells were treated to induce persistence as explained above, they were treated with 2 µg/mL doxycycline for 72 hours needed to induce about 70% of NFE2L2 silencing. Alternatively, HCT116 cells were infected with specific shRNA targeting CD44 (TRCN0000057564; Merck), SLC7A11 (TRCN0000043126; Merck) and GRIN1 (TRCN0000063369; Merck), selected by puromycin and then treated with FOLFIRI as mentioned above.

### In bulk RNA-seq

Total RNA was extracted by using Qiagen RNeasy® Mini Kit and concentration and quality was evaluated by Nanodrop 8000 spectrophotometer (Thermofisher scientific). mRNA purification and NGS libraries were obtained following Illumina instruction (TruSeq RNA Sample Preparation) and sequenced on Illumina Platform (Paired End/2×100; 5 million reads for each sample). Read tags were analysed with kallisto^82^ v0.50.1, utilizing gencode v45^83^ as the gene model. Gene-level transcript summaries were generated using txImport^84^. Differential expression was assessed through linear models in edgeR^85^.

### Enrichment analysis

Enrichment analysis was performed considering only significantly differentially expressed genes (DEGs) with an FDR < 0.01. The tool used is the Database for Annotation, Visualization, and Integrated Discovery (DAVID)^86^. Enriched pathways derived from Gene Ontology (GO) databases. The results returned from the analysis were filtered to consider only those with a p-value < 0.05.

### Statistical analysis

Data are represented as mean ± SD (if not diversely indicated). Comparisons between two groups were assessed by using two-tailed Student’s t-test. Comparison between more groups were assessed by using the one-way ANOVA followed by Dunnett post-test or Tukey post-test for multiple comparisons, as indicated in figure legend. P ≤ 0.05 and lower were considered significant. **** p ≤ 0.0001; *** p ≤ 0.001; ** p ≤ 0.01; * p ≤ 0.05.

### Data availability

Bulk RNA-seq and single cell RNA-seq analysis were performed on data previously deposited at the following accession numbers in the ArrayExpress: E-MTAB-12948, E-MTAB-12949, E-MTAB-12950, E-MTAB-12951. Data have been processed as previously reported^1^.

The data sets generated in this study for RNA-seq studies have been deposited at the following accession number E-MTAB-16489. Correspondence and requests for materials should be addressed to Giovanni Tonon and Simona Punzi.

## Acknowledgments

We thank all the members of the Tonon laboratory for the discussions, support and for critical reading of the manuscript. We are grateful to the Flow cytometry Resource, Advanced Cytometry Technical Applications Laboratory (FRACTAL), the Electron Microscopy BioImaging Centre (ALEMBIC) and the Experimental Imaging Centre (CIS) at Ospedale San Raffaele IRCCS. Graphical abstract was created with BioRender.com. This work was supported by Fondazione AIRC (AIRC, Investigator Grant # 28987 (to G.T.)). S.P. was supported by the grant from the Italian Ministry of Education, University and Research, project PNC0000001 D34 Health—Digital Driven Diagnostics, prognostics and therapeutics for sustainable Health care “CUP” B53C22006090001; G.G. was supported by the Italian Ministry of Health grant (RF-2021–12374586); C.F. was supported by the grant from the European Union - Next Generation EU - NRRP M6C2 - Investment 2.1 Enhancement and strengthening of biomedical research in the NHS” PNRR-MCNT1-2023-12378347 “CUP” C43C24000220007.

## Author contributions

S.P. and I.V. performed *in vitro* treatments on HCT116; S.P. and Gi.C. performed *in vitro* treatment on SW780 and U2OS; G.C. and E.G. assisted with the experiments; G.C., G.F.M.G., O.A.B. and C.F. performed experiments on organoids; I.V. performed cell transduction and prepared cells for RNA sequencing; S.P. and I.V. analysed experimental data and applied statistics; D.C. analysed RNA sequencing and single cell sequencing data; S.P. and G.G. performed flow cytometry and imaging experiments; M.P. and N.A. performed deep-learning analysis on organoids revised by V.B.; E.T. and A.N. provided reagents and protocols; A.N. and L.S. revised the manuscript; S.P. and G.T. conceived the project and designed the experiments; S.P. and G.T. wrote the manuscript with input from all authors.

## Ethic

Investigations have been conducted in accordance with ethical standards and according to national and international guidelines. Human Tissue biopsies were collected from patients whose informed consent was obtained in writing according to the policies of the Ethics Committee of the San Raffaele Hospital and the regulations of the Italian Ministry of Health. The studies were conducted in full compliance with the Declaration of Helsinki.

**Correspondence** and requests for materials should be addressed to Simona Punzi and Giovanni Tonon.

## Supplementary material

### Supplementary Figures

**Suppl. Fig.1.**
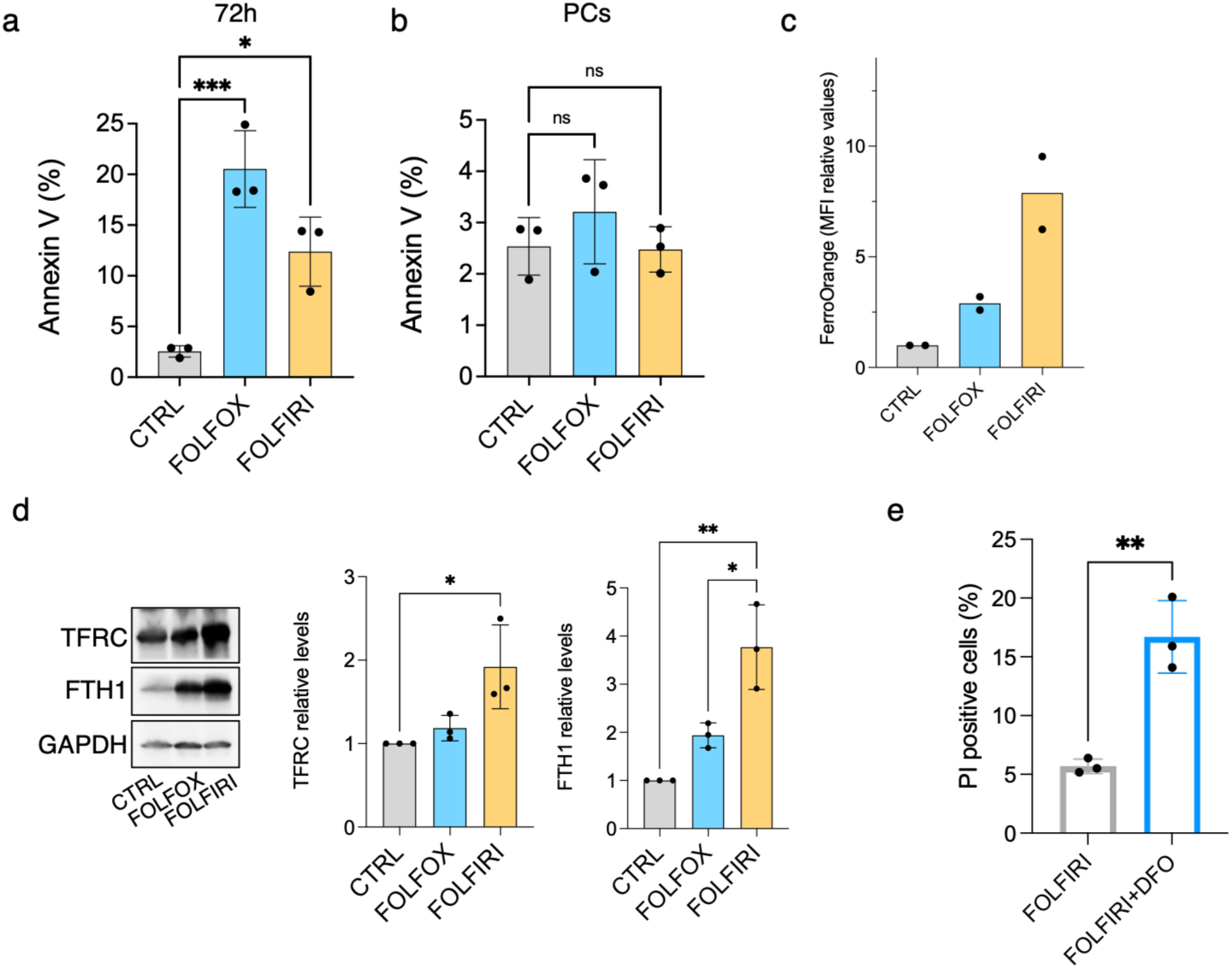
CRC PCs inhibit programmed cell death mechanisms to survive chemotherapy. **a-b.** Annexin V positivity expressed as percentage (%) of positive cells quantified at flow cytometry evaluated upon 72h of FOLFOX or FOLFIRI treatments (**a**) or in PCs upon 2 weeks administration (**b**). Significant differences among groups calculated by ANOVA followed by Dunnett post-test (n=3 biological replicates; mean±SD; *:P<0.05; ***: P<0.001; ns: not significant). **c.** Labile iron content in PCs and parental CTRL quantified by FerroOrange assay by flow cytometry analysis (n=2 biological replicates; mean). **d.** Western blot analysis and respective relative quantification of TFRC and FTH1 protein levels in parental HCT116 CTRL and PCs. GAPDH was used as a normalizer. Significant differences among groups calculated by ANOVA followed by Dunnett post-test (n=3 biological replicates; mean±SD; *:P<0.05; **: P<0.01). **e.** Propidium iodide positive cells in FOLFIRI-PCs treated with DFO. Significant differences calculated by Student’s t-test (n=3 biological replicates; mean±SD; **: P<0.01).

**Suppl. Fig.2.**
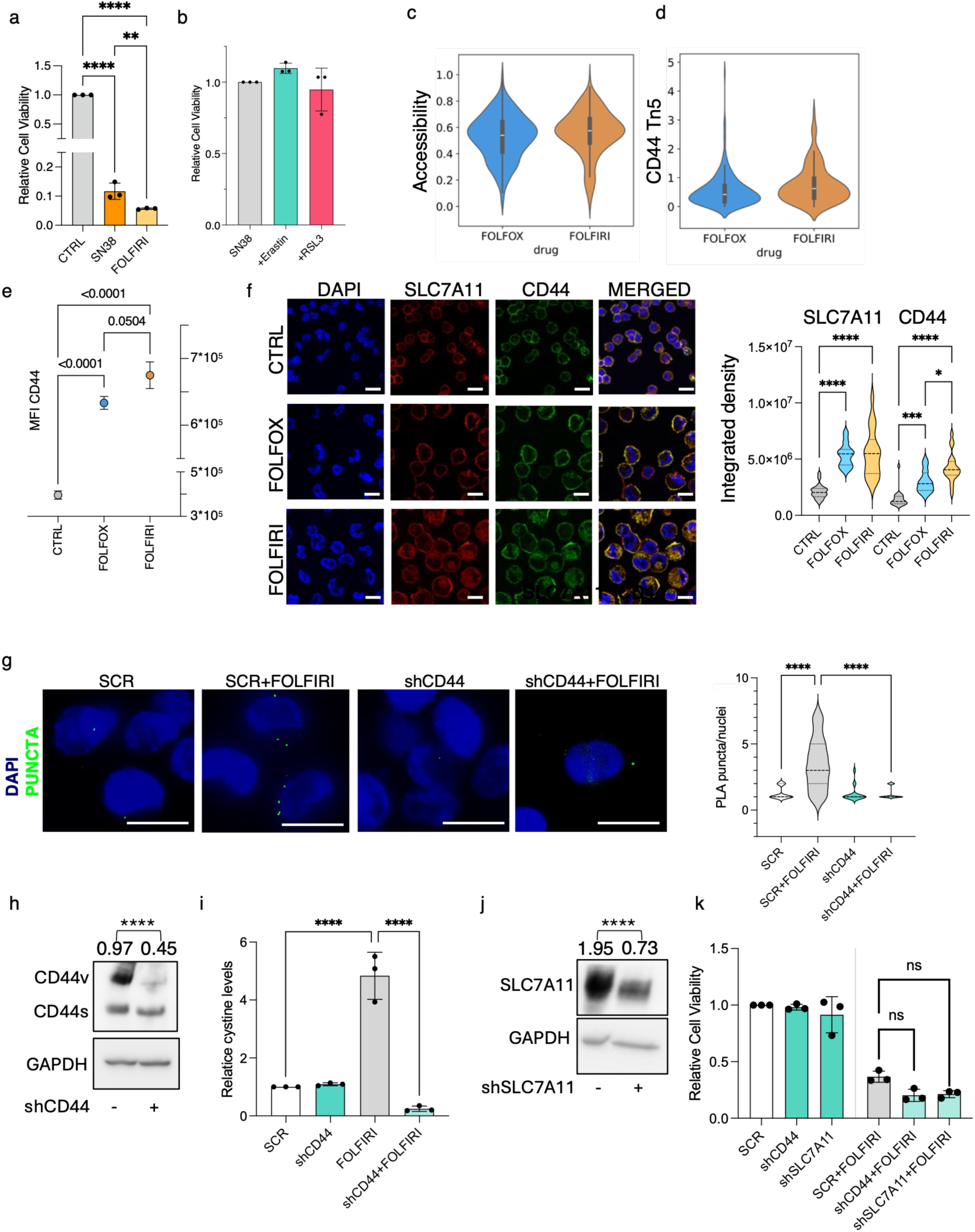
FOLFIRI-PCs activate Xc^-^ antiporter system. **a.** Relative cell viability of HCT116 cells treated with SN38 or FOLFIRI for 72h. Significant differences among groups calculated by ANOVA followed by Tukey post-test (n=3 biological replicates; mean±SD; **: P<0.01; ****: P<0.0001). **b.** Relative cell viability of PCs obtained upon two weeks of SN38 treatment and Erastin or RSL3 administration. Significant differences among groups calculated by ANOVA followed by Dunnett post-test (n=3 biological replicates; mean±SD). **c-d.** Single cell Get-sequencing analysis of global chromatin accessibility state (**c)** in PCs or related to tn5 of CD44 specific regions (**d**). Differences among the two groups were calculated by Student t-test (FOLFOX vs FOLFIRI - c: P=0.00325; d: P= 0.00415). **e.** Flow cytometry analysis of CD44 levels in CTRL, FOLFOX- or FOLFIRI-PCs. Significant differences among groups calculated by ANOVA followed by Tukey post-test (n=3 biological replicates; mean±SD). **f.** Immunofluorescence analysis of CD44 and SLC7A11 in CTRL, FOLFOX- or FOLFIRI-PCs. Significant differences among groups calculated by ANOVA followed by Tukey post-test (n=3 biological replicates biological replicas and 20 cells analysed; mean±SD; *: P<0.05; ***: P<0.001; ****: P<0.0001). **g.** Proximity Ligation Assay (PLA) foci quantified as PLA puncta/nuclei in cells treated with FOLFIRI or after CD44 silencing (scale bar: 10 µm). Significant differences among group calculated by ANOVA followed by Tukey post-test (n=3 biological replicates biological replicas; 10 cells per condition: ****: P<0.0001). **h**. Representative western blot of CD44 levels in control or shCD44 cells. Significant differences were calculated on density levels of CD44v on GAPDH, used as normalizer (mean of n=3 biological replicates biological replicas; Student t-test; ****: P<0.0001). **i.** Relative cystine uptake in FOLFIRI and CD44 silenced cells (shCD44). Significant differences among groups calculated by ANOVA followed by Tukey post-test (n=3 biological replicates; mean±SD; ****: P<0.0001). **j.** Representative western blot of SLC7A11 levels in control or shSLC7A11 cells. Significant differences were calculated on density levels of SLC7A11 on GAPDH, used as normalizer (mean of n=3 biological replicates biological replicas; Student t-test; ****: P<0.0001). **k.** Relative cell viability of FOLFIRI treated cells infected to silence CD44 or SLC7A11. Significant differences among groups calculated by ANOVA followed by Tukey post-test (n=3 biological replicates; mean±SD; ns: not significant).

**Suppl. Figure 3.**
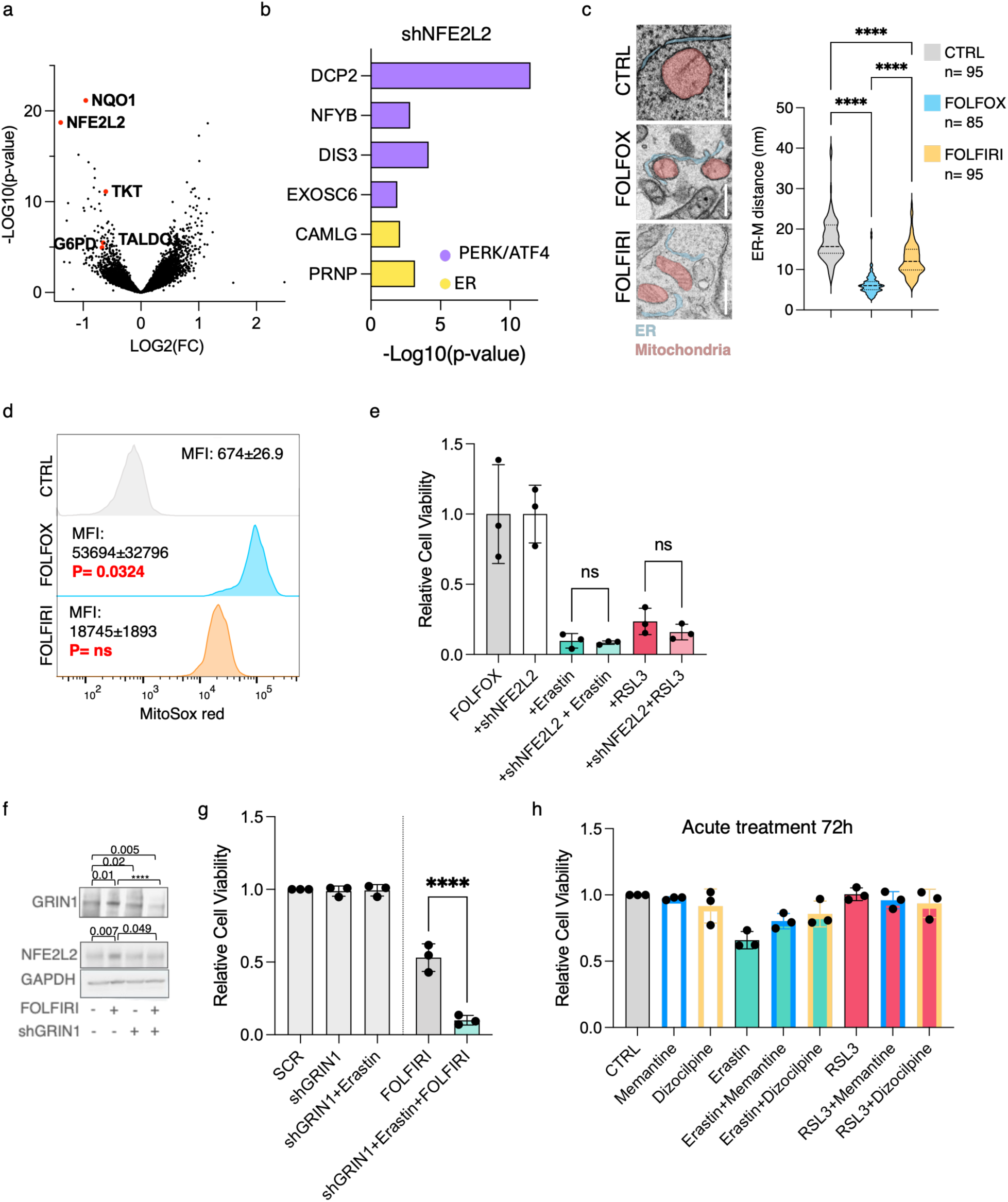
NMDAR-NFE2L2 axis inhibition resensitizes PCs to ferroptosis. **a-b.** Volcano plot representing LOG2 fold change (FC) and corresponding -LOG10(p-value) of genes upon NFE2L2 genetic silencing (shNFE2L2). Known downregulated direct targets (**a**) or upregulated genes related to PERK/ATF4, or ER (**b**) were highlighted. **c.** Representative TEM images and corresponding quantification of mitochondria/endoplasmic reticulum distance expressed in nm. Scale bar: 500nm. Significant differences among groups calculated by ANOVA followed by Tukey post-test (median, first and third quartile are shown; ****: P<0.0001). **d.** Representative histograms and corresponding values quantifying mitochondrial ROS content (MFI; n=3 biological replicates; mean±SD) by flow cytometry analysis. Statistical differences among CTRL and PCs were calculated by applying an ANOVA followed by Dunnett post-test. **e.** Relative cell viability of FOLFOX-PCs in shLuc or shNFE2L2 conditions treated with Erastin or RSL3. Significant differences among groups calculated by ANOVA followed by Tukey post-test (n=3 biological replicates; mean±SD; ns: not significant). **f.** Representative western blot of NFE2L2 and GRIN1 levels in cells treated with FOLFIRI or in shGRIN1 condition. Significant differences were calculated on density levels of NFE2L2 or GRIN1 on GAPDH, used as normalizer (mean n=3 biological replicates biological replicas; Student t-test; ****: P<0.0001). **g.** Relative cell viability of FOLFIRI-PCs in shGRIN1 conditions treated with Erastin. Significant differences among groups calculated by ANOVA followed by Tukey post-test (n=3 biological replicates; mean±SD; ****: P<0.0001). **h.** Relative cell viability of HCT116 parental cells treated with Memantine or Dizocilpine, in presence or Erastin or RSL3.

**Suppl. Fig. 4:**
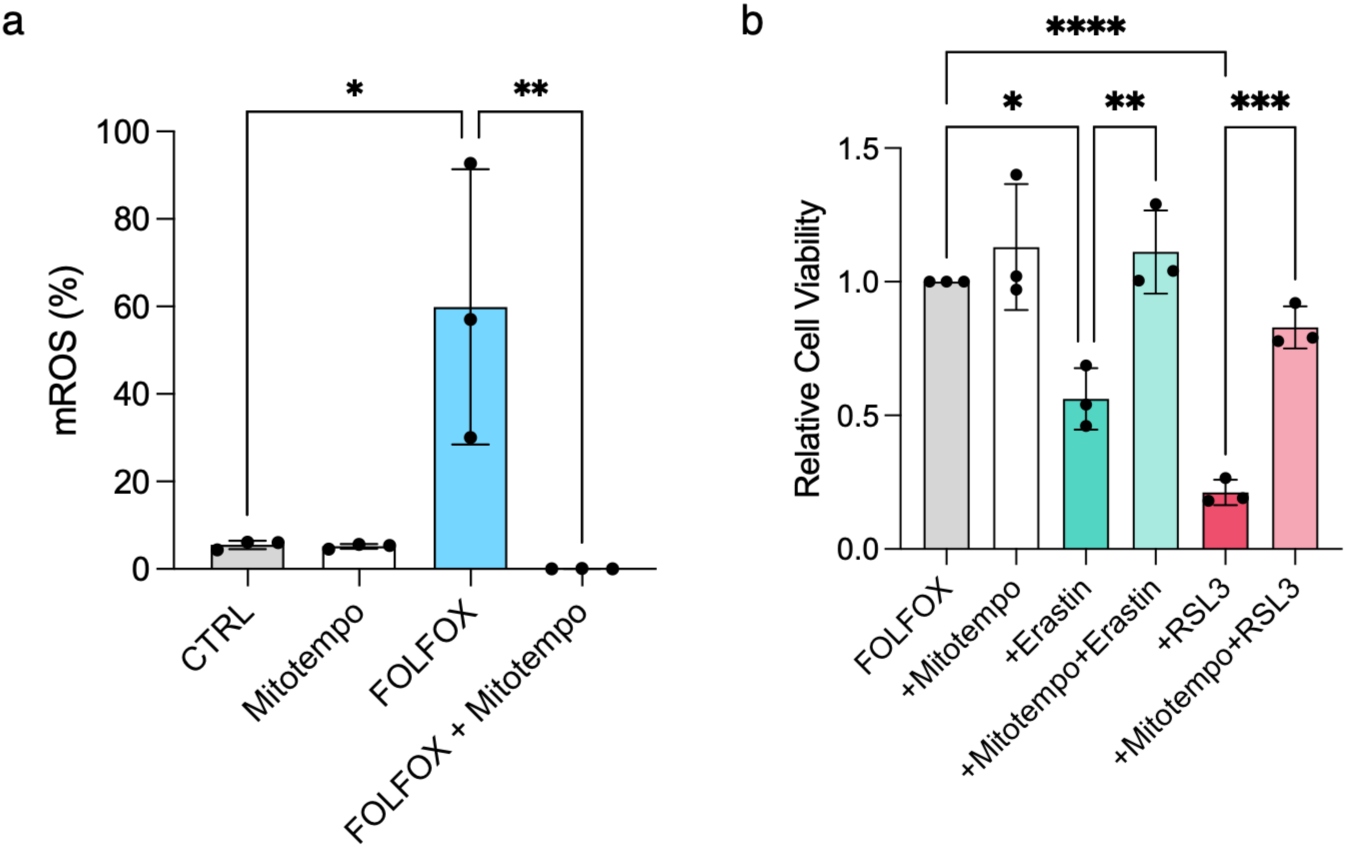
mROS drive ferroptosis in PCs. **a.** Mitochondrial ROS quantified in FOLFOX-PCs and after Mitotempo treatment detected by flow cytometry. Significant differences among groups calculated by ANOVA followed by Tukey post-test (n=3 biological replicates; mean±SD; *:P<0.05; **: P<0.01). **b.** Relative cell viability of FOLFOX-PCs treated with ferroptosis inducers Erastin and RSL3 and Mitotempo. Significant differences among groups calculated by ANOVA followed by Tukey post-test (n=3 biological replicates; mean±SD; *: P<0.05; **: P<0.01; ***: P<0.001; ****: P<0.0001)

**Suppl. Fig. 5.**
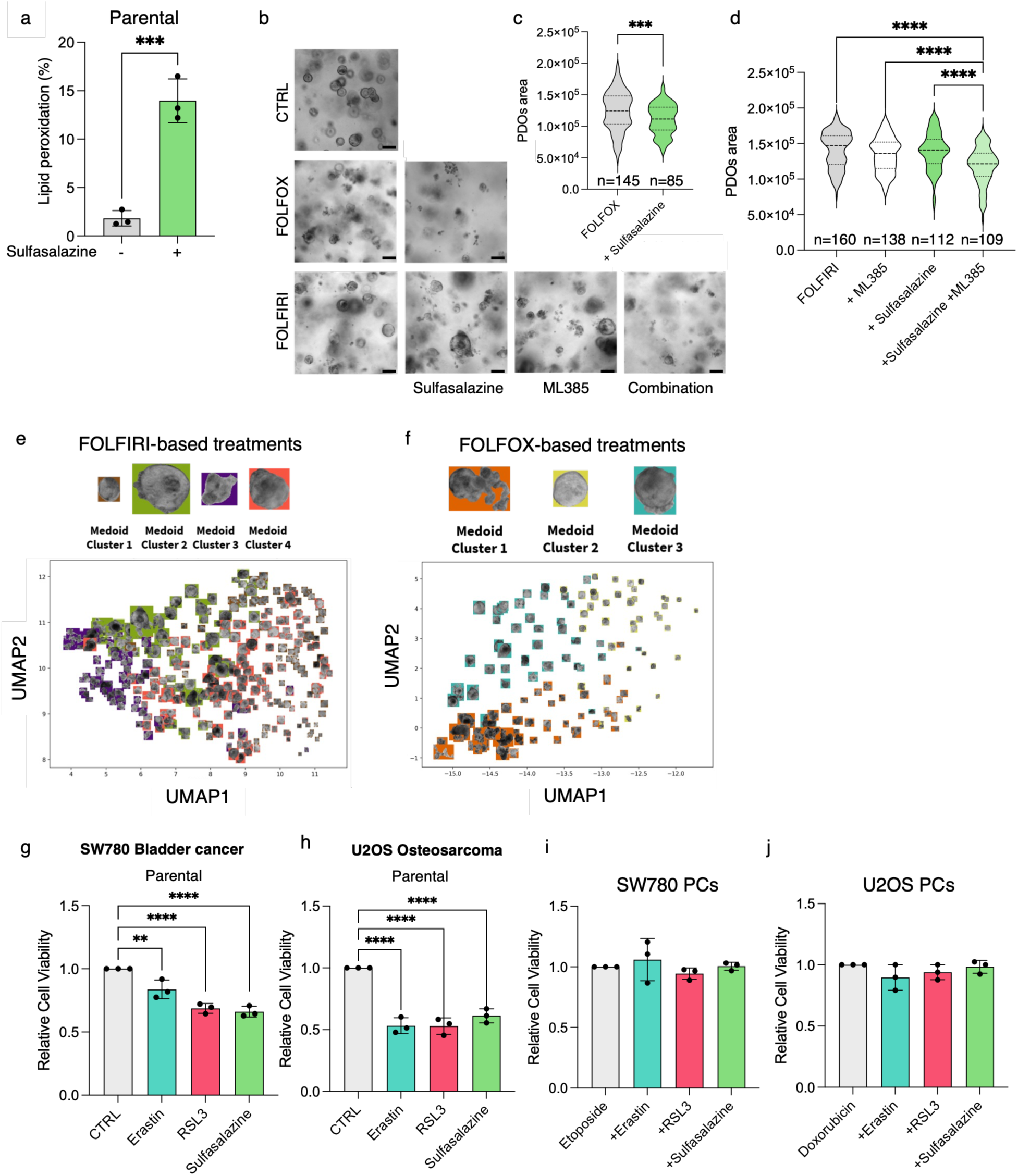
Topoisomerase inhibition in several cancer models. **a.** Lipid peroxidation levels quantified as percentage positivity (%) in parental HCT116 upon Sulfasalazine treatment by flow cytometry analysis. Significant differences among groups calculated by two-tailed Student’s t-test (n=3 biological replicates; mean±SD; ***: P<0.001). **b.** Representative images of PDOs treated with chemotherapies, sulfasalazine, ML385 and their combination. Scale bar: 100 µm. **c.** Persister PDOs viability upon treatment with FOLFOX and sulfasalazine. Significant differences among groups calculated by two-tailed Student’s t-test (mean±SD; ***: P<0.001). **d.** Persister PDOs viability upon treatment with FOLFIRI, sulfasalazine and ML385. Significant differences among groups calculated by ANOVA followed by Tukey post-test (mean±SD; ****: P<0.0001). **e-f.** Cluster medoids obtained by the global feature-architecture of PDOs upon FOLFIRI- (**e**) or FOLFOX- (**f**) based treatments. UMAP projection of labeled instance embeddings and representative cluster medoids. **g-j.** Relative cell viability of parental (**g-h**) or persisters (**i-j**) SW780 bladder cancer and U2OS osteosarcoma cells treated with Erastin, RSL3 or Sulfasalazine for 72h. Significant differences among groups calculated by ANOVA followed by Dunnett post-test (n=3 biological replicates; mean±SD; **: P<0.01; ****: P<0.0001).

### Supplementary Table

**Suppl. Table 1.** Reactome analysis of 708 upregulated genes in FOLFIRI-PCs. P-value <0.05.

